# fluxTrAM: Integration of tracer-based metabolomics data into atomically resolved genome-scale metabolic networks for metabolic flux analysis

**DOI:** 10.1101/2024.11.26.625485

**Authors:** Luojiao Huang, German Preciat, Jesus Alarcon-Gil, Edinson L. Moreno, Agnieszka Wegrzyn, Ines Thiele, Emma L. Schymanski, Amy Harms, Ronan M.T. Fleming, Thomas Hankemeier

**Affiliations:** Metabolomics and Analytics Centre, Leiden Academic Centre for Drug Research, Leiden University, Leiden, Netherlands; Computer Sciences Department, University of Guadalajara, Guadalajara, Mexico; Department of Neurobiology, Care Sciences and Society, Division of Neurogeriatrics, Center for Alzheimer Research, Karolinska Institutet, Stockholm, Sweden; Digital Metabolic Twin Centre, School of Medicine, University of Galway, University Rd, Galway, Ireland; School of Medicine, University of Galway, University Rd, Galway, Ireland; Luxembourg Centre for Systems Biomedicine, University of Luxembourg, Belvaux, Luxembourg

## Abstract

Quantitative inference of intracellular reaction rates is essential for characterising metabolic phenotypes. The classical experimental method for measuring metabolic fluxes makes use of stable-isotope tracing of metabolites through the metabolic network, followed by mass spectrometry analysis. The most common ^13^C-based metabolic flux analysis requires multidisciplinary knowledge in analytical chemistry, cell biology, and mathematical modelling, as well as the use of multiple independent tools for handling mass spectrometry data. Besides, flux analysis is usually carried out within a small network to validate a specific biological hypothesis. To overcome interdisciplinary barriers and extend flux interpretation towards a genome-scale level, we developed fluxTrAM, a semi-automated pipeline for processing tracer-based metabolomics data and integrating it with atomically resolved genome-scale metabolic networks to enable flux predictions at genome-scale. fluxTrAM integrates different software packages inside and outside of the COBRA Toolbox v3.4 for the generation of metabolite structure and reaction databases for a genome-scale model, labelled mass spectrometry data processing into standardised mass isotopologue distribution data (MID), and metabolic flux analysis. To demonstrate the utility of this pipeline, we generated ^13^C-labeled metabolomics data on an *in vitro* human induced pluripotent stem cell (iPSC)-derived dopaminergic neuronal culture and processed ^13^C-labeled MID datasets. In parallel, we generated a cheminformatic database of standardised and context-specific metabolite structures, and atom-mapped reactions for a genome-scale dopaminergic neuronal metabolic model. MID data could be exported into established flux inference software for conventional flux inference on a core model scale. It could also be integrated into the atomically resolved metabolic model for flux inference at genome-scale using moiety fluxomics method. The core model flux solution and moiety flux solution were then compared to two additional flux solutions predicted via flux balance analysis and entropic flux balance analysis. The extensive computational flux analysis and comparison helped to better evaluate the obtained flux feasibility of the neuron-specific genome-scale model and suggested new tracer-based metabolomics experiments with novel labeling configurations, such as labelling a moiety within the thymidine metabolite. Overall, fluxTrAM enables the automation of labelled liquid chromatography (LC)-mass spectrometry (MS) data processing into MID datasets and atom mapping for any given genome-scale metabolic model. It contributes to the standardisation and high throughput of metabolic flux analysis at genome-scale.

## Introduction

Fluxomics aims for quantification of reaction flux at a genome-scale to comprehensively characterise cellular metabolism. Quantification of reaction flux is challenging as, unlike molecular species abundance, intracellular fluxes cannot be directly measured but must be indirectly inferred from measurements of labelled molecular species [1]. A typical in vitro fluxomic workflow proceeds from experimental design and cell culture implementation, to mass spectrometric data generation and processing, and finally to inference of reaction flux by computational modelling [2]. Therefore, progress in fluxomics is dependent on overcoming challenges in a set of complementary fields, including cell biology, analytical chemistry, and mathematical modelling. Herein, we focus on overcoming challenges in automated data handling, processing of mass spectrometry data and preparing an atom mapped model, which ultimately enables computational inference of metabolic flux.

Metabolomics allows a broad profiling of metabolites that connect diverse biochemical reactions in a metabolic network. Given a chromatographic co-elution of metabolite isotopologue peaks, reliable measurement of stable isotopically labeled metabolites requires a high capacity for analytical technology in molecule separation or peak resolution. Recent developments in mass spectrometry (MS) coupled with separation chromatography (gas chromatography, GC, or liquid chromatography, LC) improve the measurement accuracy of metabolite labelling patterns, as well as increase the coverage of metabolite classes [3,4]. In this way, metabolic flux profiling within a large-scale model becomes possible and can have the advantage of providing a systematic view of metabolism. Along with this, an advanced flux profiling approach at a genome-scale level is required.

By measuring a complex biological sample, high resolution (HR) LC-MS generates a large volume of MS raw data especially with a long analytical run for sufficient chromatographic separation. With the challenges of increased targeted metabolites, co-elution adding to the mass spectral complexity and multiple biological replicates in a tracer experiment, it becomes more difficult to process isotopologue labelling data. Manual peak integration to obtain the mass isotopologue distribution (MID) of each target metabolite becomes increasingly time-consuming and error-prone. To improve data quality and reproducibility, several automated peak detection and extraction packages have been developed for LC-HRMS [5–8], that enable automation of tracer-based metabolomics data processing. For instance, X^13^CMS provides unbiased retrieval of isotopologue groups from isotopically labelled compounds between experimental conditions [6]. MetExtact focuses on identifying the entire labelled metabolome and annotating unknown metabolites from mixtures of uniformly highly isotope-enriched and native biological samples [8]. mzMatch–ISO offers its functionality in both automated untargeted isotope annotation and relative quantification of metabolite isotopologues [7]. For our goal of targeted MID analysis, a pre-evaluation of each tool’s suitability for incorporation into a standardised workflow is indispensable.

The advancement of high-throughput experimental technology has increased the demand for computational approaches to integrate complementary sources of omics data [9–11]. One such approach is integration of omics data with genome-scale computational models of biochemical networks, which are reconstructed from experimental data on molecular species and the biochemical reactions [12]. Genome-scale models are especially useful for predicting metabolic reaction flux [13]. However, quantitative inference of intracellular metabolic reaction flux by integration of MID data with a genome-scale metabolic model has not yet been demonstrated, but rather only with a subnetwork model [14].

Integration of experimental MID data with metabolic modelling is currently challenging. On the one hand, the input of experimental MID data always needs external pre-processing including correction of naturally occurring isotopes and tracer purity, which involves intense user manipulation. On the other hand, currently available ^13^C flux analysis programs implement non-linear optimization by minimizing the difference between measured and iteratively simulated MIDs to compute an optimal flux solution within a specified metabolic network [15–18]. Adequate literature support and modelling expertise are required to formulate an organism-specific and flux consistent network model. The reconstructed model needs manually assigned atom transitions and is usually with a limited network model size. Automating the MID data processing and atom transitions resolving into a continuous workflow would be beneficial to accelerate metabolic modelling at genome-scale.

Metabolic networks are interconnected pathways of biochemical reactions within organismal system through which building blocks or compounds necessary for cellular functioning are assembled (anabolism) or energy and matter are produced by breaking down biomolecules (catabolism) [19]. A reaction in a genome-scale metabolic network can be represented by a set of atom-mappings, each of which connects an atom in a substrate metabolite to this atom in a product metabolite. Such atom-mappings can be algorithmically predicted at genome-scale [20] and when compared with manually curated biochemical reactions, accurately predict atom mapping for ∼90% of human metabolic reactions [21]. Connecting sets of atom mappings for a metabolic network enables one to atomically resolve a metabolic network at genome-scale [13]. Based on this approach, automated workflows for atom mapping have been developed for several species, including human [21] and *Arabidopsis thaliana* metabolism [22]. However, to have atomically balanced metabolic reactions, it is necessary to identify the specific metabolic structures for genome-scale models. Without this precision, atomically unbalanced reactions may result from inaccuracies in metabolite structures or incorrect reaction stoichiometry. By including external cheminformatics software such as Open Babel [23], CXCALC [24] and the Reaction Decoder Tool (RDT) [25] it is possible to convert between chemical formats, identify the pH required to balance the number of hydrogens in the substrates and products, and thereby atom-mapped balanced metabolic reactions.

Graph theoretical analysis of an atomically resolved metabolic network enables identification of conserved moieties, each of which is a set of atoms that remains invariant with respect to all metabolic transformations in a given network [26]. The existence of conserved moieties means that a biochemical network is a special type of hypergraph [26], in the sense that the underlying stoichiometric matrix has special mathematical properties not shared with a general rectangular matrix corresponding to a general hypergraph. Fundamentally, this special property arises because one can represent a biochemical network at an atomic level [27]. In a companion paper [28], we exploited this property to develop a new mathematical and computational method that linearly relates metabolic reaction flux to the rate at which each labelled or unlabelled conserved moiety transitions between metabolites. This *moiety fluxomics* method enables inference of metabolic reaction flux at genome-scale, given mass isotopologue distribution data.

Herein, we present flux inference by integrating stable-isotope laeblled metabolomics data with atomically resolved genome-scale model (fluxTrAM), as an extension to the COBRA Toolbox v3.4 [13]. Without reinventing the wheel, fluxTrAM took advantage of existing third-party packages to achieve automated atom mappings and labelled LC-MS data processing. It generates a cheminformatic database of standardised and context-specific metabolite structures based on their InChI string and atom-mapped reactions for a given genome-scale metabolic reconstruction. In parallel, wrapper function interface to third-party packages, to enable processing of LC-MS raw data by fluxTrAM and then generate standardised MID data as labelling measurement input, which is subsequently used for various approaches to flux inference. The MID and atom mapping data obtained can be combined to estimate the internal fluxes at genome-scale [28]. The utility of the pipeline was demonstrated by using stable-isotope labelled LC-MS data from a human dopaminergic neuronal culture. fluxTrAM integrated tracer-based metabolomics data with genome-scale flux analysis and compared with conventional ^13^C metabolic flux analysis. Based on the genome-flux solution, the pipeline also predicted which labelled moiety to be considered for labelling when designing a new labelling experiment to further explore dopaminergic neuron metabolism.

## Materials and methods

An overview of the methodology is given in **Figure 1** and described in detail below.

**Figure 1.**
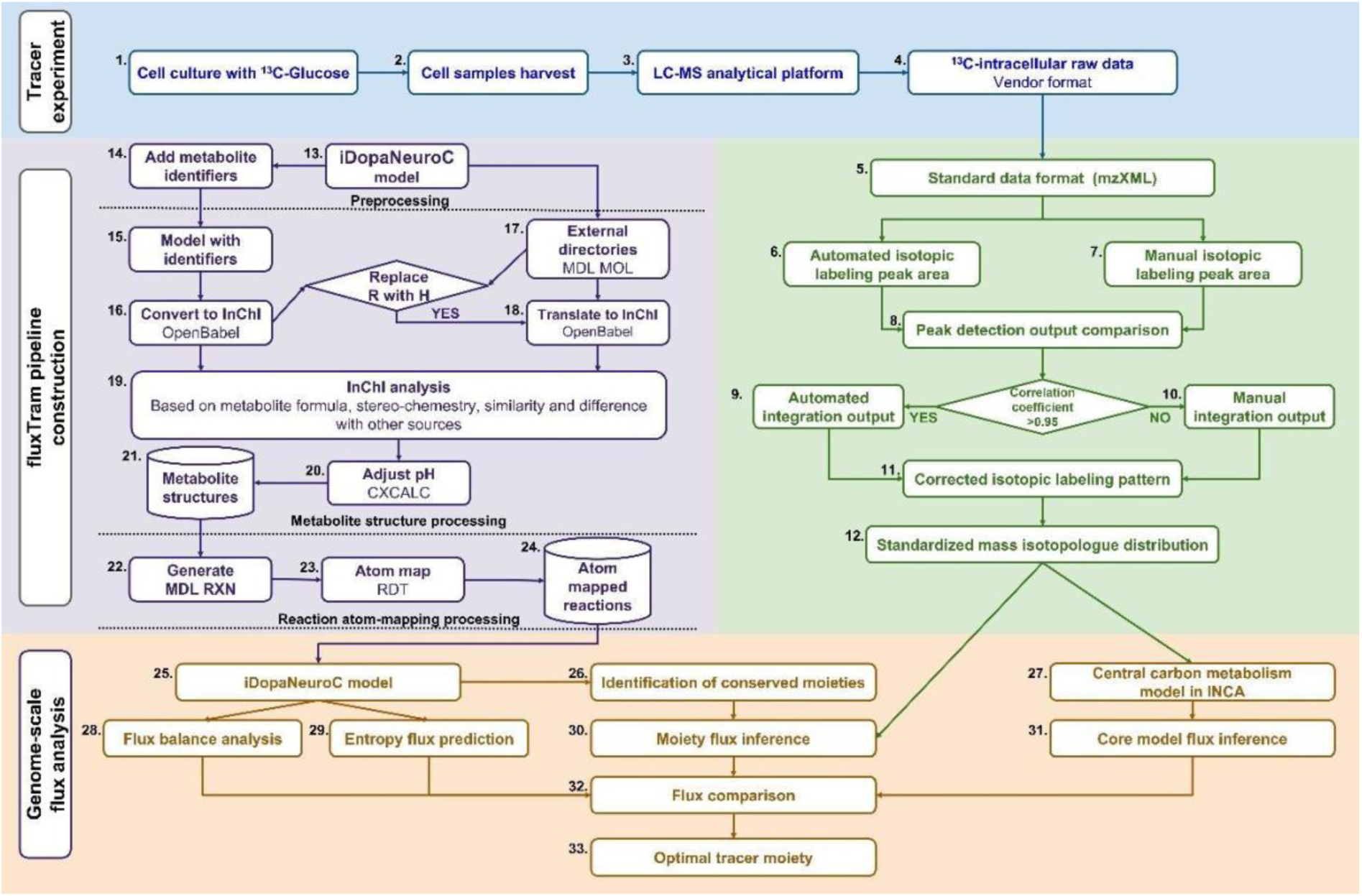
Methodological overview. *In vitro* cell culture was fed with fully ^13^C labelled glucose (1). Labelled cellular samples were collected after certain exposure time (2). A validated LC-MS platform [29] was established to measure labelled intracellular samples (3). Labelled intracellular raw mass spectral data were converted into a standard format (4, 5) and automated peak integration was used to obtain an isotopic labelling pattern (6, 9), unless it showed low correlation (8) to that obtained by manual peak integration (7, 10). Isotopic labelling patterns were converted to mass isotopologue distribution data in a standard format (11, 12). In parallel, atom mapping data was obtained from a genome-scale metabolic network by adding structural cheminformatic data for each metabolite in the network if required (13-15), or using existing external data (17). The structural cheminformatic data for each source in the model was used to generate an InChI (16, 18) where any R group in a metabolite structure was replaced by a hydrogen atom. The InChI strings obtained were compared to select the most representative structure for the genome-scale model (19). The number of hydrogen atoms of the highest-scoring metabolite structures was adjusted to match the number of hydrogens at which the metabolite is represented in the metabolic model of dopaminergic neuronal metabolism (iDopaNeuroC) (20), resulting in a database of standardised metabolite structures (21). The metabolic network stoichiometry and metabolite structure database were used to express each reaction in an atomically resolved reaction format (22), which was then atom-mapped (23) to generate an atom-mapped reaction database (24). Atom mapping data was used to identify the conserved moieties in the iDopaNeuroC model (25, 26). The standardised mass isotopologue distributions together with conserved moieties were used as constraints on the iDopaNeuroC model to infer fluxes using the moiety fluxomic method (30). Fluxes inferred using the moiety fluxomic method were compared with predictions of fluxes without any isotopologue constraints on the iDopaNeuroC using flux balance analysis (FBA, 28) and entropic flux balance analysis [30] (29). Furthermore, the standardised mass isotopologue distribution data was imported established flux inference software (INCA [16,31]) using a central carbon metabolism model (27, 31). We compared inferred fluxes generated from the four different approaches: moiety flux inference, FBA flux prediction, entropy flux prediction and central carbon metabolism flux inference using INCA (32). Optimal genome-scale flux solutions were selected and used to further screen candidate tracer moieties for a new labelling culture experiment (33).

## 1. Tracer experiment

### 1.1 In vitro cell culture

Generation of an *in vitro* culture of midbrain-specific dopaminergic neurons followed an established protocol [32,33], with the adaptions described below. This culture method was the same as that used for generation of iDopaNeuroC model [30] a context-specific model of dopaminergic neuronal metabolism.

### N2B27 medium preparation

The culture medium, denoted N2B27 medium, was used as the basis to prepare both maintenance and differentiation media. 49.25 mL of culture medium was obtained by mixing 24 mL Neurobasal medium (Invitrogen/Life Technologies), 24 mL of Dulbecco’s modified Eagle’s medium (DMEM)/F12 medium (Invitrogen/Life Technologies) supplemented with 1% penicillin and streptomycin (Life Technologies), 0.5 mL of 200 mM L-glutamine (Life Technologies), 0.5 mL of B27 supplement without Vitamin A (Life Technologies) and 0.25 mL of N2 supplement (Life Technologies). Glucose-free N2B27 medium was made in the same way, but with the Neurobasal medium replaced with Neurobasal medium with no D-glucose (Invitrogen/Life Technologies), and the DMEM/F12 medium replaced with stable isotope labeling with amino acids in cell culture (SILAC) advanced DMEM/F-12 Flex medium with no glucose (Invitrogen/Life Technologies).

### Plate coating

Cell-culture treated 12-well plates (ThermoFisher scientific) were coated with 1% Matrigel (Discovery Labware, Inc., USA, Catalogue number 354277) in 600 μL of DMEM (1X) medium supplemented with knockout serum replacement (ThermoFisher scientific).

### Cell seeding and maintenance

At the time of cell seeding, the knockout DMEM (1X) medium from the coating step was removed from each well and the K7 hNESC line was seeded in three replicate wells. The medium to maintain the hNESC in culture, denoted maintenance medium, was based on N2B27 medium with 0.5 μM PMA (Enzo life sciences), 3 μM CHIR (Axon Medchem) and 150 μM ascorbic acid (Sigma Aldrich). The cell seeding was done by preparing 1.8×10e6 million cells/mL in maintenance medium and adding 300 μL of this preparation together with another 300 μL of maintenance medium to reach 4×10e5 cells per well. The plate was incubated at 37 °C and 5% CO2 for 48 h.

### Neuronal differentiation and maturation

The differentiation medium with PMA was prepared to induce the differentiation of hNESC towards midbrain dopaminergic neurons and consisted of N2B27 medium with 200 μM ascorbic acid, 0.01 ng/μL BDNF (Peprotech), 0.01 ng/μL GDNF (Peprotech), 0.001 ng/μL TGFβ3 (Peprotech), 2.5 μM dbcAMP (Sigma Aldrich) and 1 μM PMA. This medium was completely replaced every 2 days during the next 6 days of culture in the differentiation process. For the maturation of differentiated neurons, PMA is required to be absent from the differentiation medium [32,33]. This differentiation medium without PMA was used from day 9 onwards and complete media replacement was done every 2 days for 2 weeks.

### ^13^C-labelled neuron culture and sample collection

^13^C-labelled differentiation medium without PMA was prepared using glucose-free N2B27 medium, supplemented with 20.4 mM fully carbon labelled U-^13^C6-glucose (Cambridge Isotope Laboratories, USA), 200 μM ascorbic acid, 0.01 ng/ μL BDNF, 0.01 ng/ μL GDNF, 0.001 ng/ μL TGFβ3, 2.5 μM dbcAMP. For the pilot stable isotopic labelling study set, differentiated neurons were maintained in two groups separately with ^13^C labelled and unlabelled medium. Three replicate wells of neurons from each group were incubated under 37 °C and 5% CO2 condition for 4 hours before quenching. For the formal stable isotopic labelling study set, dopaminergic neurons were cultured in labelled medium at 37 °C and 5% CO2 conditions with different incubation times, from 0min, 5min, 10min, 20min, 40min, 1h, 2h, 3h, 5h, 8h, 12h to 24h. Each condition group was run in triplicates. The spent medium was collected into a 1.5 mL Eppendorf tube. Neurons in each well were immediately quenched by using ice-cold 80% MeOH in water and harvested into a 1.5 mL Eppendorf tube as cell lysate. All samples were fast frozen into liquid nitrogen and stored in the -80 °C freezer until measurement.

### 1.2 Sample measurement

Labelled cellular samples were measured on a liquid chromatography-mass spectrometry (LC-MS) platform to obtain the mass isotopologue distribution (MID) of target metabolites (*SM4. Table S1*) [29]. The concentrations of U-^13^C6-glucose in cell and medium were quantified using the same platform. The cell lysate was sonicated, vortexed and then centrifuged at 16000 relative centrifugal force (rcf), at 4 ℃ for 10 min. Cell pellets were collected to measure the protein content using a bicinchoninic acid (BCA) assay (Thermo Fisher Scientific Inc, USA). Supernatants were transferred into clean 1.5 mL Eppendorf tubes and evaporated to dryness in a Labcono SpeedVac (MO, United State). Each sample was reconstituted with 60 μL ice cold methanol/water (80%/20%; v/v). 50 μL of the reconstitution volume was collected and transferred into a new Eppendorf tube as a cell supernatant sample. Next, 50 μL (5 μL) of each cell supernatant (medium) sample was treated with liquid-liquid extraction by adding 40 μL of ice-cold methanol/water (80%/20%; v/v), 45 μL of ice cold milliQ water and 65 μL of ice-cold chloroform, followed with mixing and vortexing for 5 min and centrifuging at 16000 rcf 4 ℃ for 10 min. 130 μL of the aqueous phase was transferred into a new Eppendorf tube and extracted again by adding 25 μL of ice-cold methanol/water (50%/50%; v/v) and 65 μL of ice-cold chloroform, followed with mixing and vortexing for 5 min and centrifuging at 16000 rcf, 4 ℃ for 10 min. 140 μL of the upper aqueous phase was collected and taken to dryness. The residue was reconstituted with 50 μL of methanol/water (50%/50%; v/v), vortex-mixed, and centrifuged to remove debris. The supernatants were finally transferred to vials for LC-MS analysis.

LC-MS analysis was performed on a SCIEX tripleTOF 5600 MS system (SCIEX, USA) coupled to a Waters Acquity UPLC Class II (Waters, USA) equipped with a SeQuant® ZIC®- cHILIC HPLC column (2.1 mm x 100 mm, 3.0 μm, Merck, Germany). The LC-MS method was previously reported [29]. Mobile phase A was 90% acetonitrile, 10% water with 5 mM ammonium formate, and mobile phase B was 10% acetonitrile, 90% water with 5 mM ammonium formate. LC elution followed a nonlinear gradient that starts from 0% B (0-2 min) to 40% B (20 min), ends up with 0% B (20.1-35 min). The flow rate was 0.25 mL/min and the injection volume was 3 μ L. The MS detection was set for a scan range of 50-900 *m/z*. Acquired raw LC-MS data files were stored in vendor-specific formats (.wiff and .wiff.scan).

## 2. Tracer-based metabolomics data processing

A pipeline for processing of tracer-based metabolomic data was developed, which takes raw mass spectrometry data as input and ultimately generates mass isotopologue distribution data in standard format as output (**Figure 2**). This pipeline is implemented in MATLAB (Mathworks Inc, USA), but calls several established external software tools for (i) LC-MS raw data conversion, (ii) Peak detection and extraction, (iii) Isotopologue peak correction, (iv) Isotopologue peak summary.

**Figure 2.**
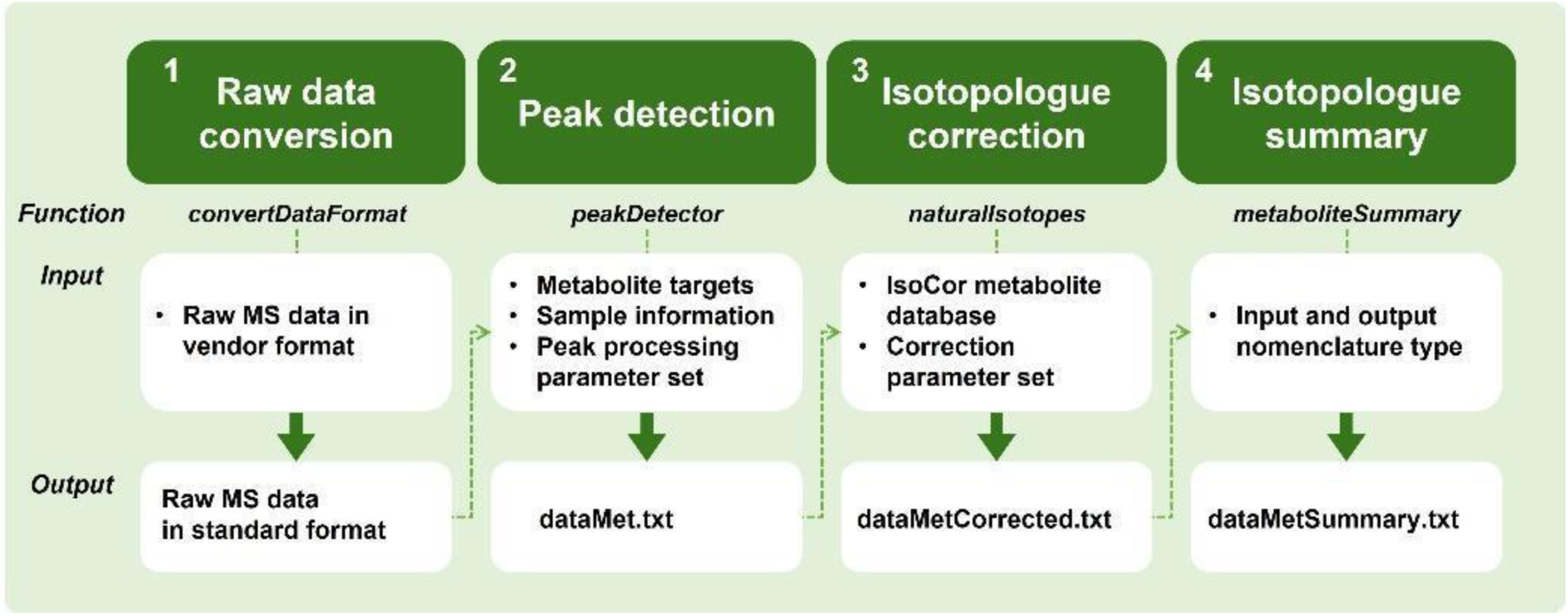
The schematic representation of tracer-based metabolomics data processing pipeline. In step 1, ’Raw data conversion’, raw LC-MS data was processed by the function ’*convertDataFormat*’ into MS data in standard format. In step 2, ’Peak detection’, standardized MS data used additional inputs for running function ’*peakDetector*’ to generate a result table of metabolite integration in a text file ’dataMet.txt’. At step 3, ’Isotopologue correction’, the dataMet.txt file, the isoCor metabolite database, and the correction parameter set were used as inputs for running the function *’naturalIsotopes*’ to remove the isotopic impurity of tracer and the naturally occurring isotopic abundance. This created a ’dataMetCorrected.txt’ file with a corrected integration table. In step 4, ’’Isotopologue summary’, the dataMetCorrected.txt file was used as an input together with the entered nonmenclature types for running function ’*metaboliteSummary*’, which finally generated a ’dataMetSummary.txt’ file with an integration table including interconverted metabolite nonmenclatures and summarized MID results shown in mean and standardized deviation.

### LC-MS raw data conversion

The function ***convertDataFormat*** was created based on ProteoWizard msconvert tool [34] for the pipeline to run ’msconvert.exe’ externally and automatically but still within MATLAB environment. It is used to convert vendor-specific formats to standard formats (mzML, mzXML, MGF, MS2/CMS2/BMS2, mzIdentML). The specific output format for fluxTrAM is centroid mode in mzXML with zlib compression.

### Peak detection and extraction

To select one automated peak extraction software package, two candidate packages (mzMatch-ISO [7] and X^13^CMS [6]) were compared with two manual peak picking software packages (ElMAVEN [35], Skyline [36]). The two manual peak integration software packages (Skyline and ElMAVEN) utilised a common target metabolite list with exact molecular formula and expected retention time for peak detection. A threshold of 10 ppm was applied so that interferences from ions with close masses could be eliminated during peak extraction. In a first step of applying automated packages, XCMS [5] was used for the initial peak detection and alignment. The optimal values for the parameter settings used in XCMS, mzMatch-ISO and X^13^CMS analysis were tested in R (*as supplementary files*: X^13^CMS_src.R, mzmatch_install.R). Five types of metabolite classes from the pilot stable isotopic labelling experiment were analysed, which included three amino acids (alanine, aspartate, glutamate), three organic acids (phosphoenolpyruvate, ketoglutarate, fumarate), one sugar phosphate (fructose 1,2-biphosphate), one tripeptide (glutathione), and one nucleotide (ATP). Pearson correlation was used to evaluate the normalised integrated peak area in automated and manual ways, and finally select one package for the automated data processing.

A wrapper function (***peakDetector, MATLAB)*** was created based on (XCMS [5], mzMatch-ISO [7]) packages for the pipeline to execute R packages still within the MATLAB environment. Converted LC-MS data was first analysed using XCMS to extract peaks. All detected peak features were then processed by mzMatch-ISO for aligning, noise filtering, gap-filling peaks, and later forming into a combined PeakML file containing all samples. Based on the input of a targeted metabolite list, automated profiling could be performed to extract relevant isotopologue peaks.

### Isotopologue peak correction

For the correction of incomplete labeling of the isotopic tracer and naturally isotopic abundance in metabolites, the function ***naturalIsotopes*** was created based on IsoCor [37] for the pipeline to excute the Python module still within a MATLAB environment. Relying on the experimental inputs, such as MS resolution, isotopic tracer type, isotopic purity, the extracted isotopologue peaks for each metabolite were corrected to only preserve the isotopic abundance resulting from the isotopic tracer. The final output was the corrected MID data of each metabolite in each sample.

### Isotopologue peak summary

The function ***metaboliteSummary*** was created to summarize the MID data from each individual sample into the average MID and corresponding standard deviation. In addition, it could also recognise and convert between different possible metabolite nomenclatures (such as Virtual Metabolic Human [38] (VMH) Abbreviation or Full Name, the Kyoto Encyclopaedia of Genes and Genomes database [39] (KEGG), the Human Metabolome Database [40] (HMDB), the International Union of Pure and Applied Chemistry (IUPAC), the International Chemical Identifier (InChI) String and InChlKey [41], the Simplified Molecular-Input Line-Entry System (SMILES) [42]) through the VMH database [38].

## 3. Atomically resolving genome-scale models

A parallel novel software pipeline was developed to atomically resolve any given genome-scale metabolic model, where the result is a cheminformatic database of metabolite structures and atom mapped reactions. This pipeline is implemented in MATLAB, but calls several optional external software tools, such as Open Babel [23] and CXCALC [24] for cheminformatic data processing and the RDT [25] to atom map metabolic reactions (Java SE Development Kit is required). Installation of these external software tools is optional but recommended to obtain metabolites with higher confidence score. An overview of the methodology to generate an atomically resolved genome-scale model is given in **Figure 1** (13-24) and further described below.

### 3.1 Metabolite structures

Metabolites were represented with database identifiers, e.g., Virtual Metabolic Human [38] (VMH), PubChem [43], the Kyoto Encyclopaedia of Genes and Genomes database [39] (KEGG), Chemical Entities of Biological Interest [44] (ChEBI) or the Human Metabolome Database [40] (HMDB). In addition to database identifiers, the metabolite structures were also represented using several formats including metabolite chemical tables (MDL MOL) that list all of the atoms in a molecule, as well as their coordinates and bonds [45]; the Simplified Molecular-Input Line-Entry System (SMILES), which uses a string of ASCII characters to describe the structure of a molecule [42]; or the International Chemical Identifier (InChI) developed by the IUPAC, which provides a standard representation for encoding molecular structures using multiple layers[41] as illustrated in *Figure S2*.

For a given metabolite, comparing different metabolic databases, one may obtain substantial structural diversity due to the inclusion of different isomers in the different sources. The fluxTrAM pipeline uses all of the aforementioned databases to obtain and ultimately select a single InChI for each metabolite. InChI was chosen due to its standard structure for encoding molecular information and it is a database independent representation [41]. fluxTrAM analyses all the InChI strings obtained from each source database assigning a score based on the criteria shown in **Table 1** to identify the metabolic structure that most closely resembles the metabolite described in the genome-scale reconstruction. The InChI for each metabolite is scored considering the chemical formula excluding hydrogen atoms, similarity with other databases, stereochemistry and charge in order to avoid loss of the predictive capacity of computer models, due to propagation of an inconsistent structures.

**Table 1.** InChI based comparison. Each InChI layer contains information that can be used to identify the metabolite structure that is most similar to the metabolite in the genome-scale reconstruction.

| Concept | Score | Description |
| --- | --- | --- |
| Chemical formula | 0 or 10 | The chemical formula indicated in the genome-scale model is compared with that obtained from the InChI. This feature is given more weight in order to keep the metabolite in the genome-scale model, where all reactions are stoichiometrically consistent and mass balanced. Hydrogen atoms were ignored in this comparison since they can be modified based on the charge (**Figure S2**). |
| Charge | 0 or 1 | The charge indicated in the genome-scale model is compared with the charge obtained from the source. |
| Stereochemical information | 0 or 1 | Indicates whether the InChI contains stereochemical information or not. |
| Standard | 0 or 1 | Indicates whether the InChI is standardised or not. |
| Similarity with other databases | 0 or 1 | The number of sources where the InChI strings are identical, divided by the total number of sources. |
| Main layer similarity | 0 or 1 | The number of sources where the main layers are identical, divided by the total number of sources. |
| InChI with more layers | 0 or 1 | InChI with more layers. |

Letters in a formula that do not correspond to a chemical element, typically described with an R, represent any group of atoms attached to the rest of the molecule for example an alkyl group or hydrogen atom. To compare metabolite structures with R groups, these atoms are modified by replacing each non-chemical atom with hydrogen only for the InChI comparison (**Figure 3.1**). If the number of hydrogen atoms in the metabolite structure obtained from a database and the metabolite obtained from the genome-scale model are different, software (CXCALC [24]) is used to adjust the charge and number of hydrogen atoms of a metabolite based on the pH (**Figure 3.2**). To ensure database consistency, the molecular structures of the metabolites in the database were represented as either molecular graphs or hydrogen-suppressed molecular graphs (**Figure 3.3**). Finally, the results were presented as molecular structures in five different formats (metabolite chemical tables (MDL MOL) [45], the Simplified Molecular-Input Line-Entry System (SMILES) [42], the International Chemical Identifier (InChI), the InChIkey [41] and an image (JPEG file) containing a graphical representation of the metabolite structure (when CXCALC is installed).

**Figure 3.**
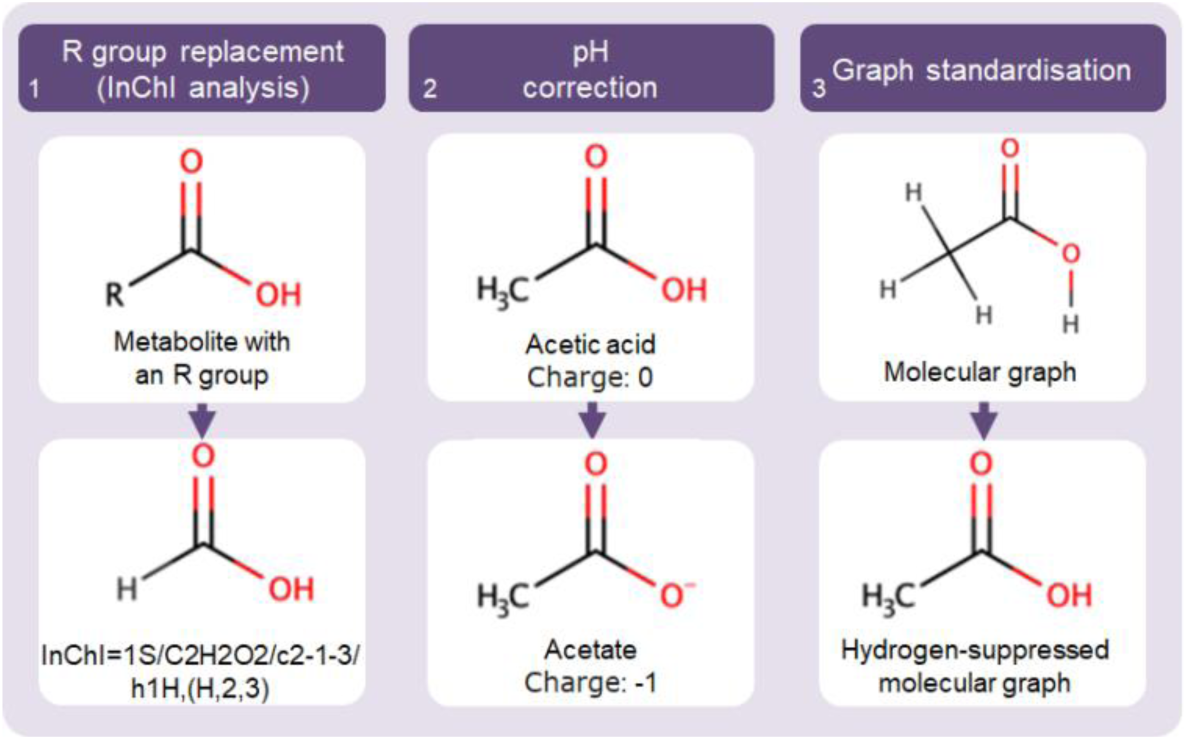
Example molecular graphs. Metabolite structure represented in different forms, as well as how it is processed by the pipeline. (1) If an unspecified alky group attached to the carboxyl group is represented by a non-chemical atom (R), the R is replaced by a hydrogen to form a molecule that can be converted into an InChI. (2) The pH of the acetic acid molecule is changed in order to obtain the charge described in the model. (3) In order to achieve implicit hydrogen standardisation, a molecular graph representing a metabolite’s structural formula is converted to a hydrogen-suppressed molecular graph, which is a molecular graph with the hydrogen vertices removed.

### 3.2 Atomically resolved reactions

A reaction database was generated for all reactions where the corresponding metabolite structures were available, as shown in the example using the metabolic reconstruction, Recon3D, which provides a comprehensive and extensive representation of human metabolism. Notably, Recon3D can also be represented as a model, featuring fewer reactions but ensuring thermodynamic, feasibility, and stoichiometric consistency; however, the reconstruction was more appropriate for this study [46] (**Figure 4**). All of these MDL mol files are available at this open source repository https://github.com/opencobra/ctf, where each filename corresponds to the VMH abbreviation for the metabolite.

**Figure 4.**
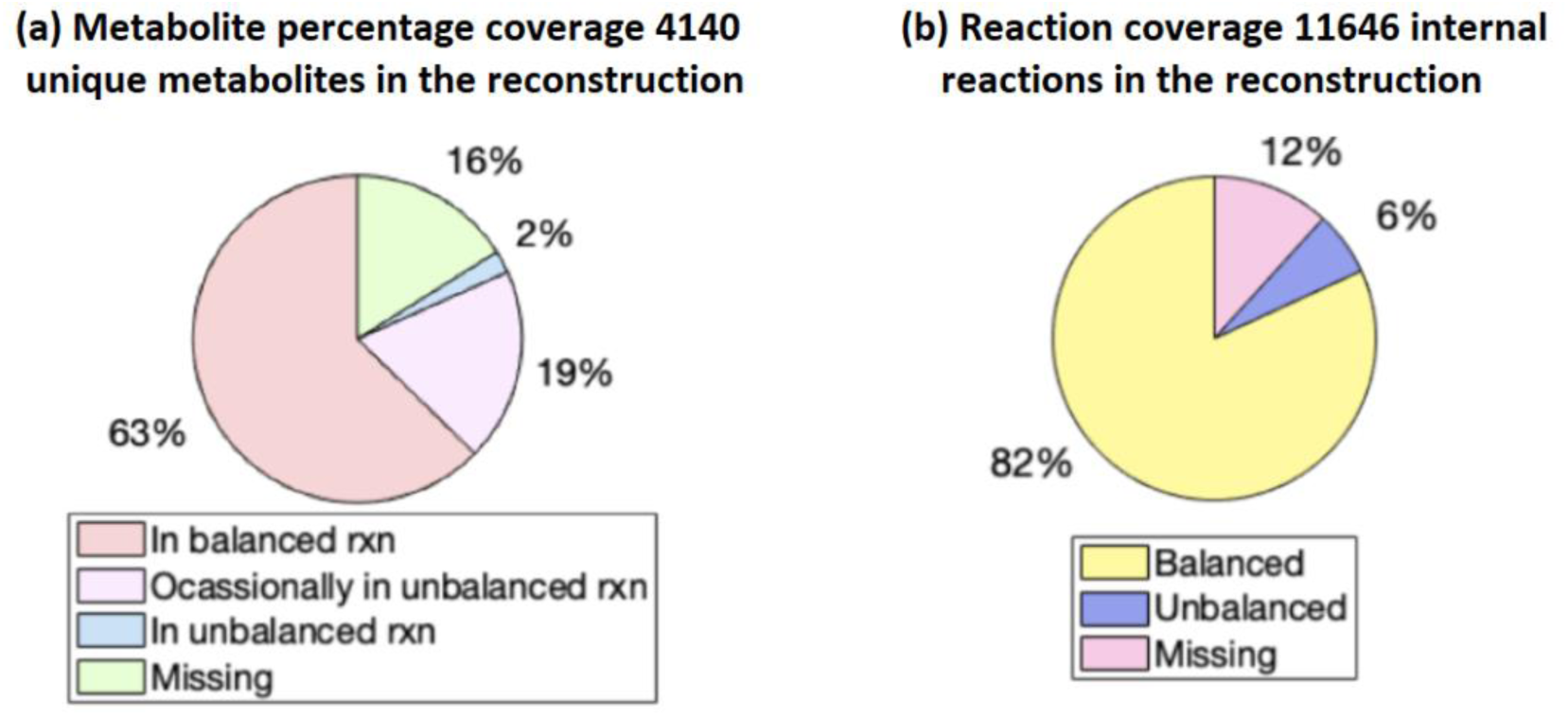
Coverage of metabolite structures and atom mapped reactions in the reconstruction case of Recon3D. (a) The fluxTrAM pipeline was used to collect 84% of the metabolite structures in Recon3D reconstruction, while only 2% correspond to the metabolite structures that are always present in atomically unbalanced reactions. (b) The metabolite structures were used to atom-map 88% of the reactions. 12% of the reactions could not be atom-mapped because at least one participating metabolite lacked structural information. Specifically, these metabolites did not have any associated database identifiers (e.g., PubChem, ChEBI) or chemical structure formats (e.g., SMILES, InChI), making it impossible to obtain their chemical structures. Additionally, 6% of the reactions were atomically unbalanced, caused by inconsistent metabolite structures.

The obtained database of metabolite structures and reaction stoichiometries from the genome-scale model was then used to generate chemical tables for each reaction (MDL RXN format) and was atom mapped using the Reaction Decoder Tool algorithm [25], interfaced with the COBRA Toolbox [13]. Atom mapped metabolic reactions were represented in reaction chemical tables (MDL RXN [45]) and reaction SMILES, while unmapped metabolic reactions were represented in International chemical identifiers for reactions [47] (RInChI; see *Figure S2*). Additionally, the reactions were atom mapped to represent their reaction mechanism. To demonstrate the general utility of this software, we generated atomically resolved reaction databases for iDopaNeuroC model [30], as well as several species-specific and community metabolic models, including the *E. coli* core model [48], models from the AGORA2 collection of 818 gut microbial metabolic models [49], and the human metabolic Recon3D reconstruction [46].

## 4. Inference of metabolic fluxes

### 4.1 Isotopomer Network Compartmental Analysis of central carbon metabolism

To perform conventional ^13^C metabolic flux analysis, a stoichiometric model of central carbon metabolism which included glycolysis, pentose phosphate pathway (PPP), tricarboxylic acid (TCA) cycle was extracted from the iDopaNeuroC model. First, this model was edited in a MATLAB-based ^13^C metabolic flux inference software package, isotopomer network compartmental analysis (INCA, [16]), describing reaction stoichiometries and carbon transitions between substrate and products (*SM4. Table S2*). Then standardised MID data was imported into INCA to infer a steady-state metabolic flux distribution in the central carbon metabolism model. INCA employs a type of decomposition of an isotopomer network to simulate MIDs as a function of metabolic flux [50], then metabolic flux is inferred by minimising the lack-of-fit between experimentally measured and computationally simulated MIDs using a Levenberg-Marquardt optimisation algorithm. Flux inference was repeated 50 times from random initial metabolic flux values to search for a global minimum fit. To assess goodness-of-fit, we subjected flux results to a chi-square statistical test and calculated 95% confidence intervals for each estimated flux value by evaluating the sensitivity of the sum of squared residuals to parameter variations.

### 4.2 Inference of metabolic flux at genome-scale with moiety fluxomics

Previously [27], we provided a mathematical definition of a conserved moiety and demonstrated how identification of the complete set of conserved moieties for a given metabolic network leads to a novel *moiety graph decomposition* of a stoichiometric matrix. We applied moiety graph decomposition to obtain the complete set of conserved moieties for a genome-scale model from human stem cell-derived, midbrain-specific, dopaminergic neurons *in vitro* [30], iDopaNeuroC. In a companion paper, we mathematically and computationally demonstrate how moiety graph decomposition enables the development of a computationally efficient constraint-based modelling method to infer metabolic fluxes from isotope labelling data for a using complete metabolic network at genome-scale. We applied this moiety fluxomic method to infer metabolic reaction fluxes at genome-scale using the aforementioned standardised MID data and genome-scale model of dopaminergic neuronal metabolism.

### 4.3 Prediction of metabolic flux at genome-scale

Two additional methods, neither of which used any tracer-based metabolomic data, were used to predict metabolic reaction flux in iDopaNeuroC, a genome-scale model of dopaminergic neuronal metabolism: (i) Flux Balance Analysis (FBA) [51], with maximisation of ATP consumption (VMHID: ATPM) to simulate a high demand for energy of dopaminergic neurons [52], (ii) Entropic Flux Balance Analysis (EFBA) maximisation of the entropy of forward and reverse fluxes, a least biased prediction given the available data [53], which also ensures that predicted flux satisfies energy conservation and the second law of thermodynamics [30]. Visualisation of all relevant flux distribution maps was performed in the software VANTED (Visualization and Analysis of Networks containing Experimental Data), v2.1.0 [54].

## Results

Aiming for quantitative flux inference at genome-scale, a semi-automated pipeline, fluxTram, was constructed, which is mainly composed of two essential parts: processing tracer-based mass spectrometry data into standardized mass isotopologue distribution; generating a cheminformatic database of standardized and context-specific metabolite structures, and atom-mapped reactions over a genome-scale model. In this study, we first evaluated the two individual modules of the fluxTrAM pipeline respectively using tracer-based metabolomics data from iPSC-derived midbrain-specific dopaminergic neuron and the corresponding iDopaNeuroC model. Next, we applied the pipeline results for conventional ^13^C flux analysis within central carbon metabolism and moiety fluxomics analysis at genome-scale, as well as flux balance analysis and entropic flux balance analysis at genome-scale.

## 1. Tracer experiment

Intracellular ^13^C-glucose increased rapidly after 5 minutes and began to stabilise within 5 hours. Meanwhile, a surge in ^13^C-glucose uptake was shown immediately at 5 min (**Figure 5**). The neuronal culture system reached a metabolic steady state after 24 hours uptake of ^13^C-glucose.

**Figure 5.**
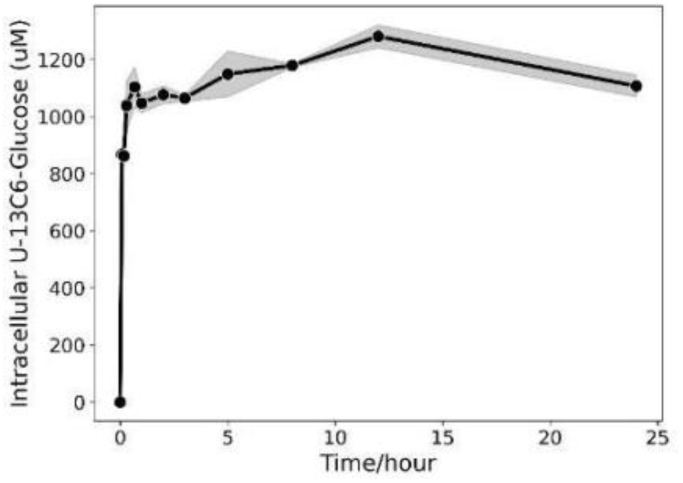
Evaluation of metabolic steady state in differentiated dopaminergic neurons by uptake of U-^13^C_6_-glucose. The intracellular level of U-^13^C_6_-glucose measured at 0min, 5min, 10min, 20min, 40min, 1h, 2h, 3h, 5h, 8h, 12h and 24h, constant level reached at 24h.

## 2. Tracer-based metabolomics data processing

### 2.1 Comparison for peak detection and extraction packages

Mass isotopologue peak integration using two automated software packages (mzMatch-ISO [7] and X^13^CMS [6]) and two manual peak picking software packages (ElMAVEN [35], Skyline [36]) were compared and the resulting isotopologue fractions for a set of 9 test metabolites are shown in **Figure 6**. The Pearson correlation of the isotopologue fractions between automated and manual is shown in **Table 2**. The test metabolites were representative of clear ion chromatographic peaks varied in peak width, intensity, and retention time. Given a list of well-established retention times and mass-to-charge ratios (*m/z*) for either respective metabolites monoisotopic peak in ElMAVEN or their mass isotopologues as well in Skyline, manual peak picking software still requires manual inspection across the integration results of every sample. Finally, ElMAVEN and Skyline showed comparable integration results for all test metabolites. The two automated packages differed in their peak integration performance. X13CMS returned isotopologue fractions that were less consistent with the integration results from the other packages. For example, no peak was extracted for phosphoenolpyruvate and compared to manual peak integration, there were low Pearson correlation coefficients for isotopologue fractions of aspartate and fructose 1,6-biphosphate. In contrast, for each of the test metabolites, mzMatch-ISO returned a similar isotopologue fraction to both of the manual software packages, which overall showed a high Pearson correlation coefficient above 0.95. Their respective statistical significance was below 0.002 (*SM4. Table S3*).

### 2.2 Peak detection and extraction evaluation

To test the peak detection and extraction performance of the incorporated mzMatch-ISO [7] package within our pipeline, we next analysed mass isotopologue distribution of essential amino acids and non-essential amino acids in neurons from the pilot study set. The integration output of 14 amino acids was compared between groups of neurons treated with labelled glucose or unlabelled glucose, the result for nine amino acids is shown in **Figure 7**. For each amino acid, the statistical differences in the isotopic incorporation between labelled and unlabelled group were evaluated based on an external t-test examination (*SM4. Table S4*). Essential amino acids can only be taken up from diet and cannot be synthesised de novo. Non-essential amino acids can be synthesised, mainly from glucose. The isotopologue fractions of six essential amino acids showed a naturally labelled isotopologue pattern in labelled neurons and had no significant difference from unlabelled neurons, including methionine, threonine, leucine, isoleucine, phenylalanine, tryptophan. In contrast, five nonessential amino acids including alanine, serine, aspartate, asparagine, glutamate showed various labelling patterns in labelled neurons. Moreover, the calculated MIDs in *Table S4* showed high consistency across the three labelled or unlabelled sample replicates. Conditionally essential amino acids, proline, tyrosine, and glutamine, showed different preferences for synthesis from glucose in neurons. A significant labelling pattern from U-^13^C6-glucose could be observed in proline. Meanwhile, glutamine and tyrosine showed little isotopic incorporation (**Figure 7**). The calculated MIDs were consistent among the labelled or unlabelled replicates.

**Figure 6.**
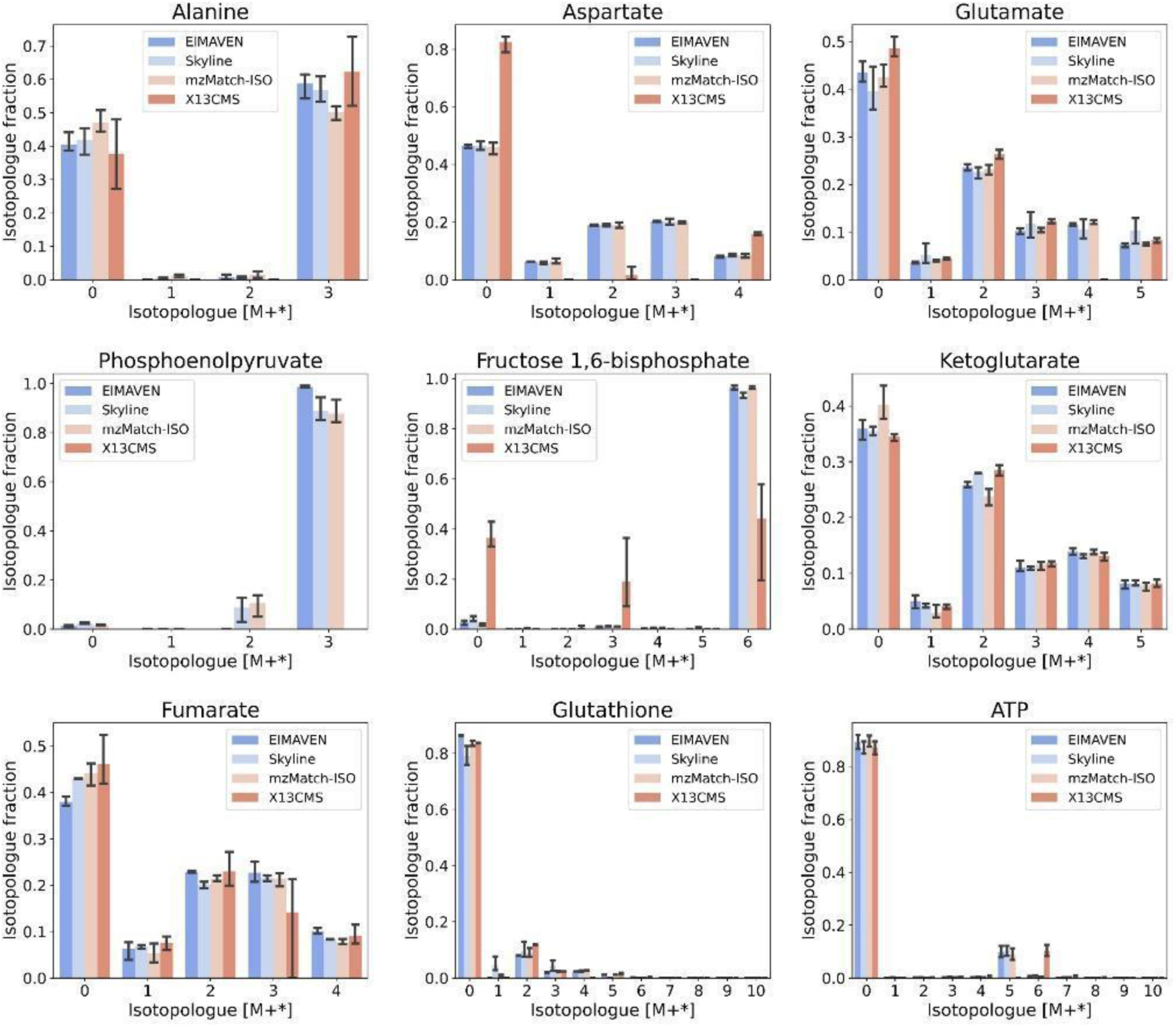
Comparison between the manual and automated peak detection and extraction results. The resulting metabolite isotopologue fractions of manual peak picking (ElMAVEN, Skyline) were compared to automated isotopologue peak integration (X^13^CMS, mzMatch-ISO) for the tested nine metabolites.

**Figure 7.**
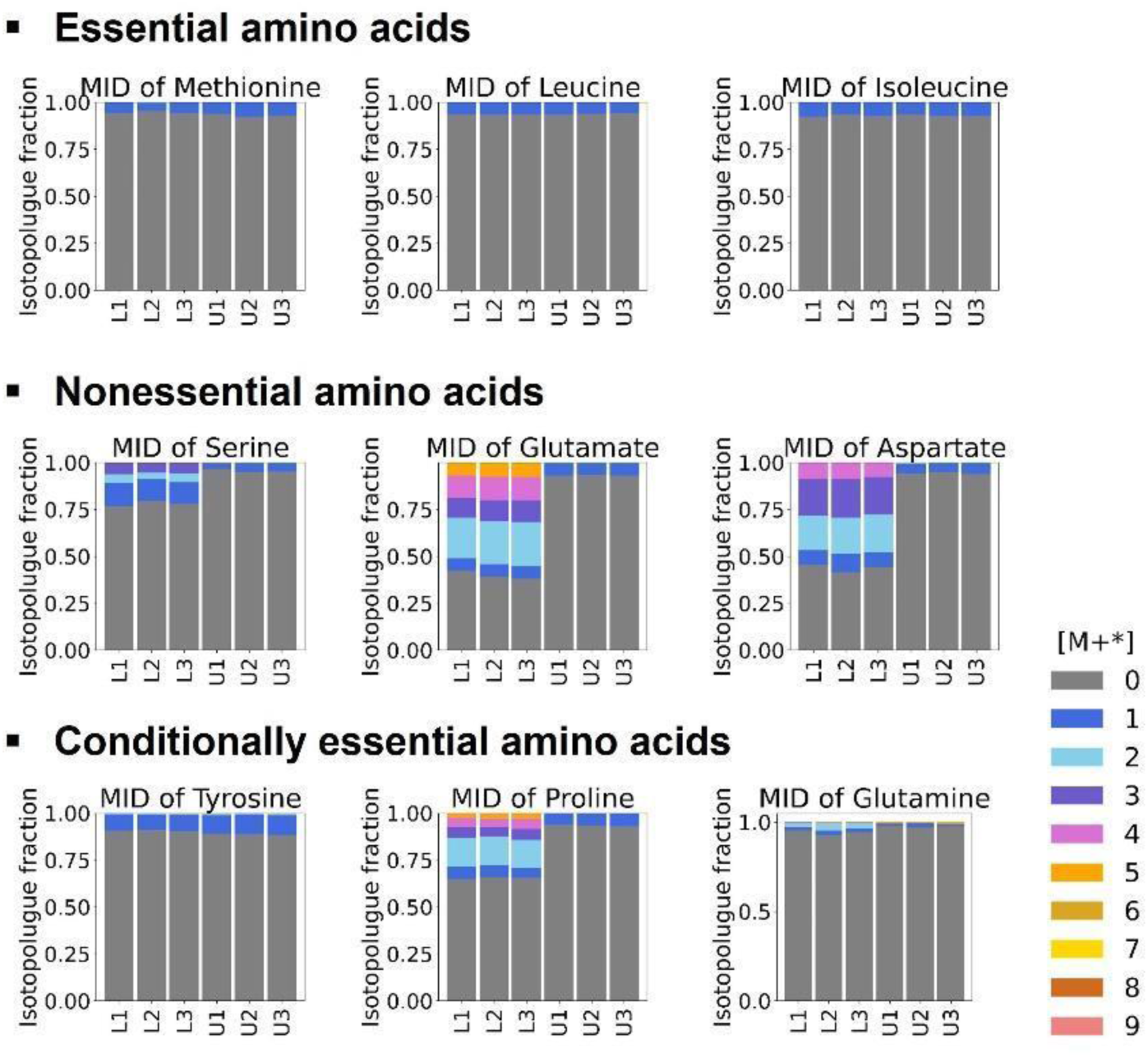
The mass isotopologue distribution (MID) of nine amino acids in the labelled neurons (n=3) and unlabelled neurons (n=3). The labelled isotopomer pattern of methionine, leucine and isoleucine (essential amino acids), serine, glutamine, aspartate (nonessential amino acids) and tyrosine, proline, glutamine (conditionally essential amino acids) shown in both labelled and unlabelled neurons. Labelled neuron samples, abbreviated in L1, L2, L3: Three replicate samples in the group of cultured neurons with isotopic tracer. Unlabelled neuron samples, abbreviated in U1, U2, U3: Three replicate samples in the group of cultured neurons with no isotopic tracer. The color represents the different detected mass isotopologues as defined by the legend.

**Table 2.** Correlation analysis of integration results from automated packages and manual software. Pearson correlation coefficients between automatically and manually collected isotopologue fractions for each of the tested nine metabolites. ND = not detected.

| Metabolite | mzMatch-ISO vs Skyline | mzMatch-ISO vs EIMAVEN | X13CMS vs Skyline | X13CMS vs EIMAVEN |
| --- | --- | --- | --- | --- |
| Alanine | 0.979 | 0.969 | 0.971 | 0.984 |
| Aspartate | 0.997 | 0.998 | 0.872 | 0.87 |
| Glutamate | 0.984 | 1 | 0.959 | 0.954 |
| Phosphoenolpyruvate | 0.99 | 0.99 | ND | ND |
| Fructose 1,6-bisphosphate | 1 | 1 | 0.645 | 0.637 |
| Ketoglutarate | 0.975 | 0.989 | 0.998 | 0.989 |
| Fumarate | 0.995 | 0.985 | 0.899 | 0.852 |
| Glutathione | 0.996 | 1 | 0.995 | 0.998 |
| ATP | 1 | 1 | 0.985 | 0.985 |

### 2.3 Data processing evaluation for a formal study set

To test the pipeline performance for processing a larger sample set, we further analysed the dynamic changes in mass isotopologue distribution of central carbon metabolites for neurons over a time course of 24 hours. A total of 36 samples for 10 metabolites (52 isotopologues) were processed automatically by fluxTram. The MID results can be found in *SM4. Table S5* and visualised in **Figure 8**. The labelling fractions of metabolites increased immediately after the initial time point. The glycolytic intermediates, phosphoenolpyruvate, pyruvate and lactate attained isotopic equilibrium within 5 hours. They showed high abundances of fully ^13^C labelled states (M+3). The tricarboxylic acid cycle, including ketoglutarate, succinate, fumarate, malate, citrate and glutamate, took nearly 24 hours to reach an isotopic steady state. Each of them showed mass isotopologue distribution in multiple ^13^C labelled states, ranging from one ^13^C labelled to fully ^13^C labelled.

**Figure 8.**
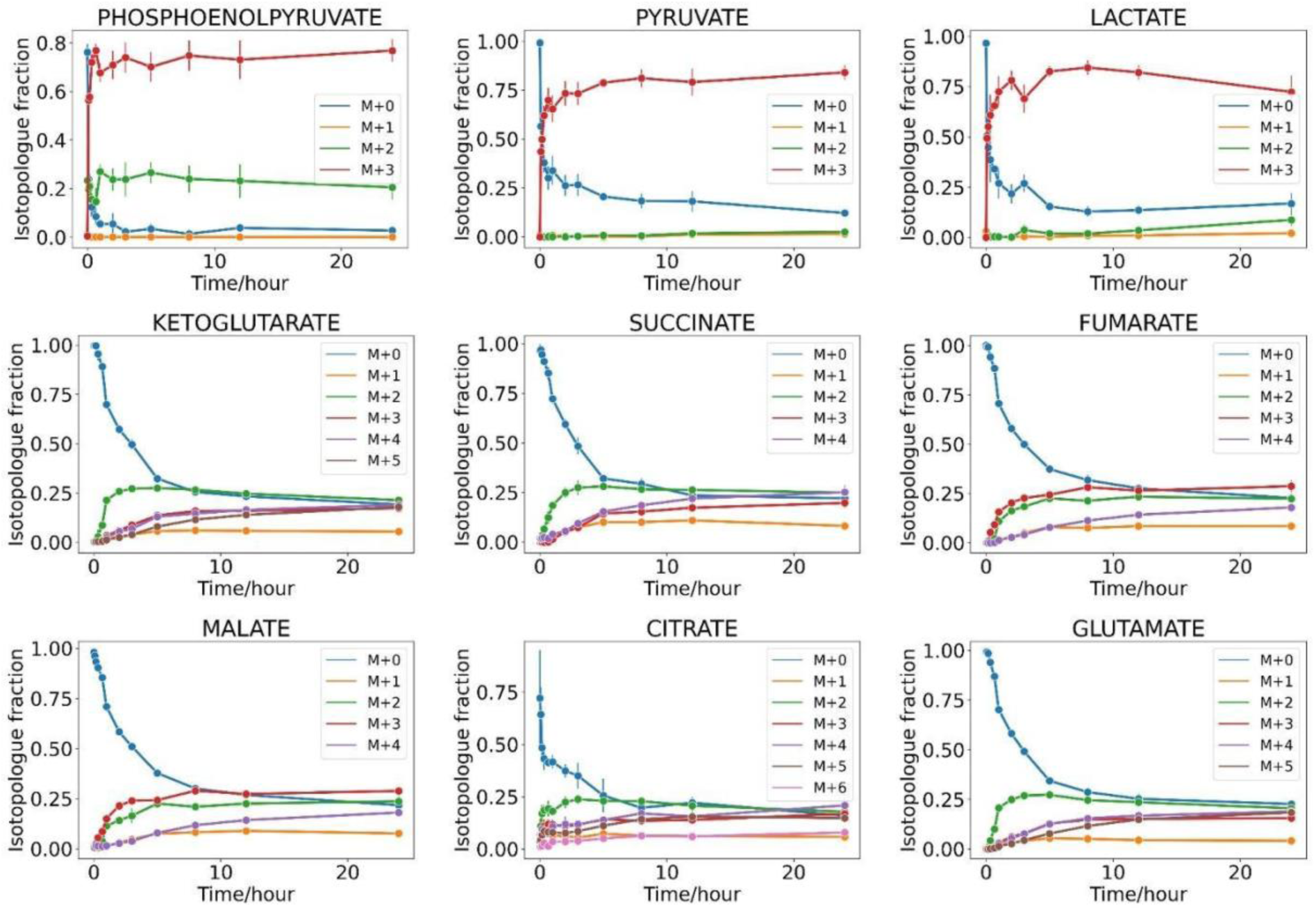
Evaluation of isotopic steady state in differentiated dopaminergic neurons by uptake of U-^13^C_6_-glucose. Mass isotopologue distributions as a function of time for intracellular metabolites within 24h of feeding isotopically labelled substrate, U-^13^C6-glucose.

## 3. Databases of metabolite structures and atom mapped-reactions

### 3.1 Metabolite structures

Figure 9 shows a comparison of the identifiers in the iDopaNeuroC model (See **Table 1**). This comparison only reflects the specificity of the metabolite identifiers in the iDopaNeuroC model, not the quality of the databases consulted. The VMH database [38] was used to refine the molecular structures in the iDopaNeuroC model, ensuring consistency with the model’s description of the chemical formula, charge, hydrogen count, and stereochemistry. While the iDopaNeuroC model is a subnetwork of Recon3D, some identifiers originally derived from external databases (e.g., KEGG, HMDB) or chemical formats (e.g., SMILES, ChEBI) did not perfectly match the described properties of the metabolites in the model. These inconsistencies, identified and resolved in the VMH database, allowed for the selection of molecular structures that most closely resemble the metabolites in the model, enabling the highest number of atomically balanced reactions. Without these corrections, reactions could remain atomically unbalanced due to mismatches in charge or hydrogen count between substrates and products. The more specific a metabolite structure is, the more likely it is to obtain a matched structure for a genome-scale metabolic network, as evidenced by Figure 9 G. The metabolite formula in the iDopaNeuroC model matched well with the formula from ChEBI [44], HMDB [40], KEGG [39], and PubChem [43] databases, however, the charge is inconsistent with the model, so they received a lower score in the InChI string analysis (Figure 9 A, B, D and E). On the other hand, the InChIs and SMILES identifiers in the model have high compatibility in the formula and charge, but they lack stereochemical information, reducing their specificity (Figure 9 C and F).

**Figure 9.**
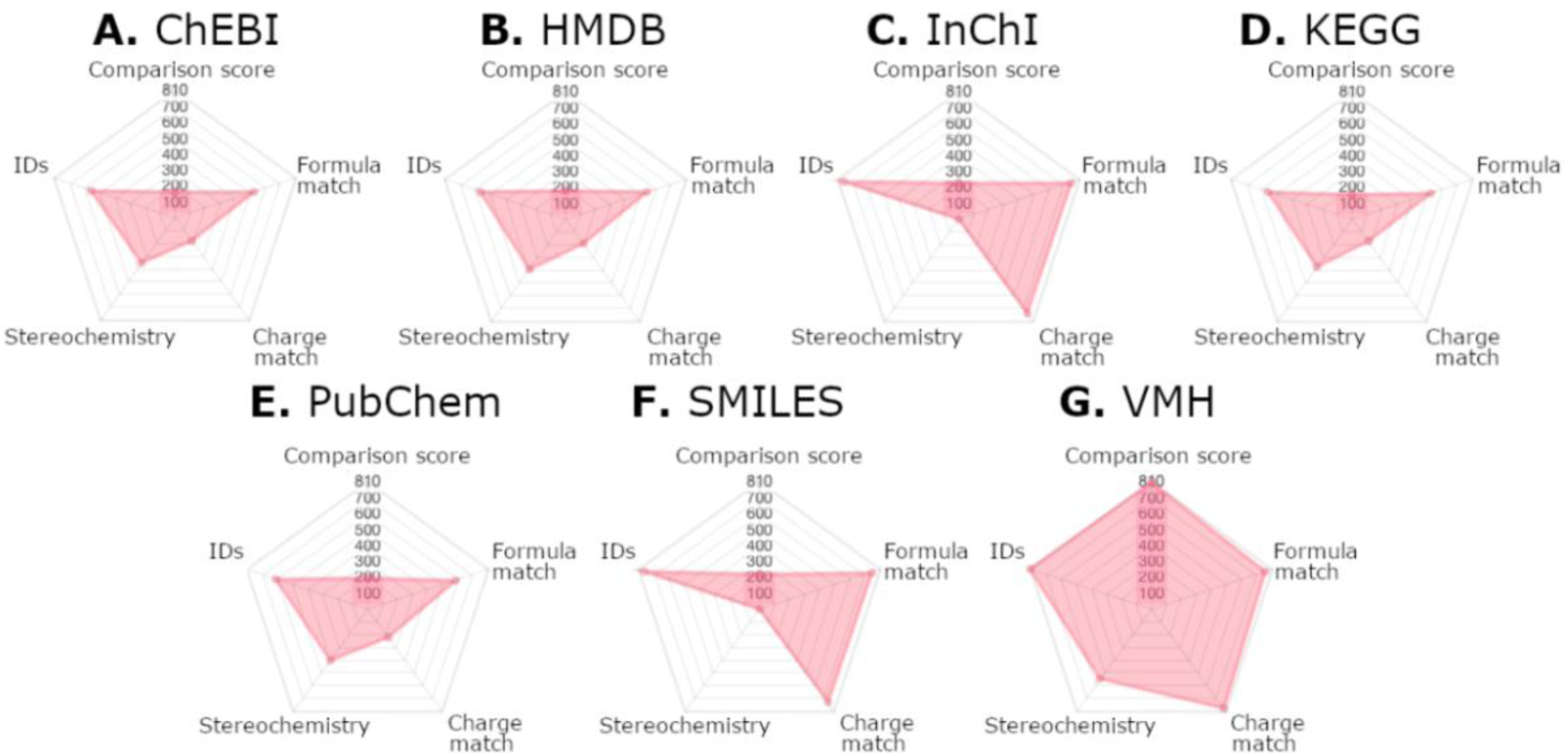
Comparison of metabolite structure identifiers in the iDopaNeuroC model. Each of the metabolite identifiers evaluated in this work, including database identifiers (e.g., ChEBI, PubChem) and chemical structure formats (InChI, SMILES), was part of the iDopaNeuroC model and represents different ways to describe a metabolite’s structure. These identifiers and formats were compared using various criteria: 1) IDs: the number of identifiers or formats evaluated; 2) Comparison score: the number of times each identifier or format received the best score based on the criteria below; 3) Formula match: the number of times the chemical formula of the identifier or format matched the metabolite formula in the iDopaNeuroC model (excluding the number of hydrogen atoms); 4) Charge match: the charge consistency of the identifier or format with the model; and 5) Stereochemistry: the number of times stereochemical information was included.

### 3.2 Atomically resolved reactions

The genome-scale model of dopaminergic neuronal metabolism, iDopaNeuroC [30], was atomically resolved. We obtained chemical structures for 810 out of 818 metabolites described in the iDopaNeuroC model. Of these, 677 metabolites were always present in atomically balanced reactions, 116 metabolites were occasionally present in atomically unbalanced reactions, and 15 metabolites were always present in atomically unbalanced reactions. Two molecular structures were not used because, while they were available, they participated in reactions were other metabolite structures were missing, rendering those reactions incomplete.

The remaining eight metabolites were categorized as "missing" because they lacked any associated database identifiers (e.g., KEGG, ChEBI) or chemical structure formats (e.g., SMILES, InChI), making it impossible to assign a molecular structure to them. These missing structures prevented the reactions involving these metabolites from being atomically resolved.

Almost all (2,254/2,262) internal metabolic reactions in the iDopaNeuroC model could be atom-mapped, of which 2,155 were atomically balanced and 99 were atomically unbalanced. Additionally, 18 metabolic reactions could not be written, because at least one metabolite structure was missing from the metabolite database. A list of the problematic reactions, along with an explanation of why each reaction was problematic, is provided in the supplementary information (SM1). **Table 3** provides a summary of the atomically resolved coverage of the iDopaNeuroC [30] model as well as several additional species and community metabolic models.

**Table 3.**
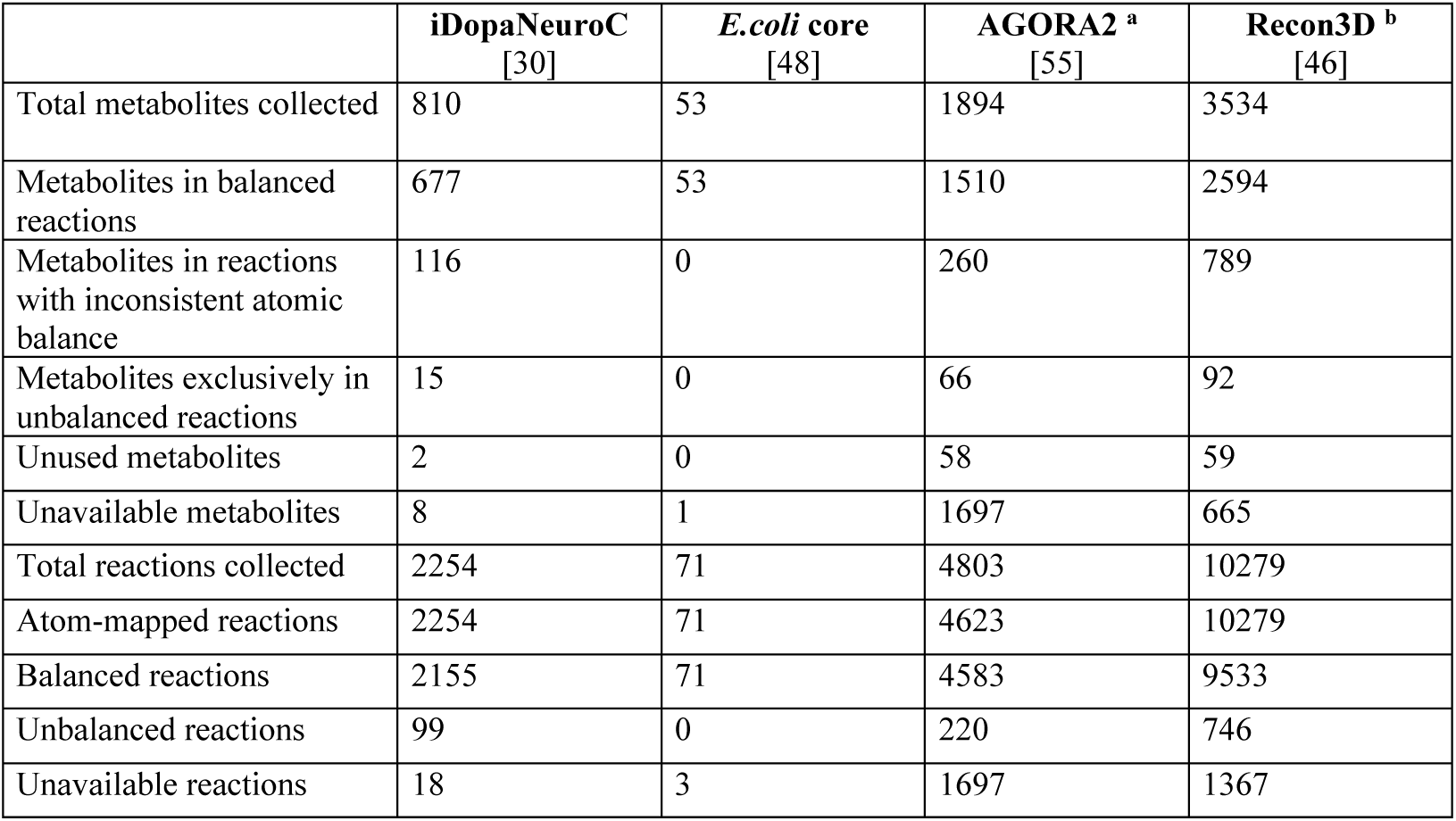
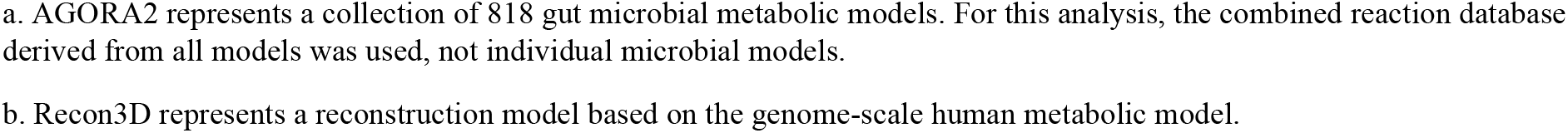
Atomically resolved coverage of metabolism in different models/reconstructions. The table summarizes the metabolite and reaction coverage in various metabolic models/reconstructions. Total metabolites collected refers to all metabolite structures gathered for the model/reconstruction. Metabolites in balanced reactions and Metabolites in unbalanced reactions reflect atomic balance in reactions. Metabolites in reactions with inconsistent atomic balance are those involved in both atomically balanced and unbalanced reactions, leading to potential structural inconsistencies, while Metabolites in exclusively unbalanced reactions only appear in reactions where atomic balance is not maintained. Unused metabolites are those structures collected but excluded from reactions due to missing structures for other participants. Unavailable metabolites lack sufficient data (e.g., chemical identifiers or formats). Total reactions collected indicates reactions successfully represented in chemical formats. Atom-mapped reactions are those containing atom-level detail. Balanced reactions and Unbalanced reactions describe atomic balance status. Unavailable reactions were not fully reconstructed due to incomplete metabolite data.

|  | iDopaNeuroC<br>[30] | *E.coli* core<br>[48] | AGORA2 <sup>a</sup><br>[55] | Recon3D <sup>b</sup><br>[46] |
| --- | --- | --- | --- | --- |
| Total metabolites collected | 810 | 53 | 1894 | 3534 |
| Metabolites in balanced reactions | 677 | 53 | 1510 | 2594 |
| Metabolites in reactions with inconsistent atomic balance | 116 | 0 | 260 | 789 |
| Metabolites exclusively in unbalanced reactions | 15 | 0 | 66 | 92 |
| Unused metabolites | 2 | 0 | 58 | 59 |
| Unavailable metabolites | 8 | 1 | 1697 | 665 |
| Total reactions collected | 2254 | 71 | 4803 | 10279 |
| Atom-mapped reactions | 2254 | 71 | 4623 | 10279 |
| Balanced reactions | 2155 | 71 | 4583 | 9533 |
| Unbalanced reactions | 99 | 0 | 220 | 746 |
| Unavailable reactions | 18 | 3 | 1697 | 1367 |

a. AGORA2 represents a collection of 818 gut microbial metabolic models. For this analysis, the combined reaction database derived from all models was used, not individual microbial models.
b. Recon3D represents a reconstruction model based on the genome-scale human metabolic model.

## 4. Inference of metabolic fluxes

### 4.1 ^13^C metabolic flux analysis of iPSC-derived midbrain-specific dopaminergic neurons

Steady-state flux inference was performed on the formal sample set collected at a 24-h time point on the basis of a stoichiometric model created in INCA covering 36 reactions (54 fluxes) from central carbon metabolism. As experimental constraints, the INCA flux analysis utilised uptake rates (glucose, glutamate), secretion rates (lactate), and ten standardised metabolite MIDs (glutamate, glutamine, ketoglutarate, succinate, fumarate, malate, citrate, lactate, pyruvate, and phosphoenolpyruvate). The final stationary 13C flux distribution results in Figure 10 showed glucose served as a primary nutrient to neurons compared to glutamine. Remarkably high glycolytic activity and relatively low metabolic activity in the tricarboxylic acid cycle were observed in neurons, whereas the pentose phosphate pathway showed almost no metabolic activity.

**Figure 10.**
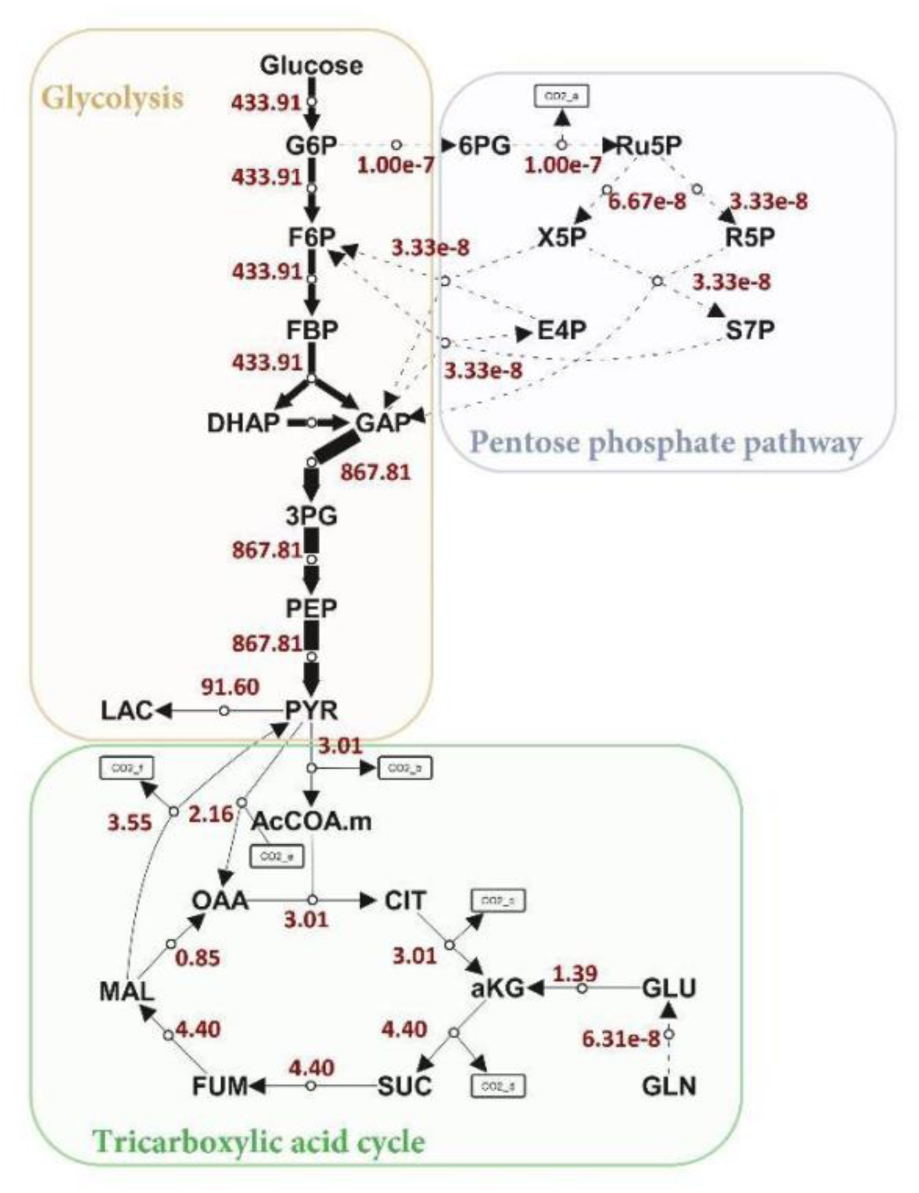
^13^C metabolic flux analysis of differentiated dopaminergic neurons within central carbon metabolism network. A steady-state flux distribution map covering glycolysis, tricarboxylic acid cycle, and pentose phosphate pathway for iPSC-derived midbrain-specific dopaminergic neurons. G6P: Glucose 6-phosphate; F6P: Fructose 6- phosphate; FBP: Fructose 1,6-bisphosphate; DHAP: Dihydroxyacetone phosphate; GAP: or G3P, Glyceraldehyde 3-phosphate; 3PG: 3-Phosphoglycerate; PEP: Phosphoenolpyruvate; PYR: Pyruvate; LAC: Lactate; AcCOA.m: Acetyl coenzyme A in mitochondria; OAA: Oxaloacetate; CIT: Citrate; aKG: alpha-Ketoglutarate; MAL: Malate; FUM: Fumarate; SUC: Succinate; GLN: Glutamine; GLU: Glutamate; 6PG: 6-Phosphogluconate; Ru5P: Ribulose 5-phosphate; R5P: Ribose 5-phosphate; X5P: Xylulose 5-phosphate; E4P: Erythrose 4-phosphate; S7P: Sedoheptulose 7-phosphate.

### 4.2 Genome-scale flux analysis

Flux balance analysis [51] and entropic flux balance analysis [30] predicted reaction fluxes in the iDopaNeuroC model without any constraints from isotope labelling data. The standardised MIDs (glutamate, glutamine, ketoglutarate, succinate, fumarate, malate, citrate, lactate, pyruvate, phosphoenolpyruvate) were used as new constraints to the iDopaNeuroC model, and helped to generate a third metabolic flux solution with a newly developed moiety solver algorithm [28]. Three genome-scale flux solutions were compared to INCA flux within the same central carbon metabolism network in order to evaluate their flux rationality.

Regarding an accurate mapping of reaction flux at the genome scale to the corresponding reaction in the core model, we adhered to the rule of combining steady-state flux, as depicted in Figure 11**.A**. The self-defined central carbon metabolism model in INCA only describes the major carbon transitions from glucose without indicating other reaction substances and cofactors. However, metabolic reactions from the iDopaNeuroC model cover multiple reactions related to the same carbon transition and with diverse reaction substances or cofactor conversions involved. For instance, phosphoenolpyruvate can use different diphosphate- nucleotides as reactants and convert into pyruvate. In this sense, we merged the net fluxes of parallel reactions that share the same carbon transitions from the iDopaNeuroC model together. In another case, a simplified reaction was described in the INCA model to represent a series of reactions in the iDopaNeuroC model. For instance, the conversion from 3-phosphoglycerate into phosphoenolpyruvate is via the intermediate metabolite 2-phosphoglycerate. Therefore, the minimum flux of consecutive reactions was used to represent the simplified reaction flux for the iDopaNeuroC model. In a third case, if the intermediate metabolite receives net fluxes both from substrates A and C, the final flux representing conversion from A to C is considered to be zero. Based on the calculated flux distributions in the same core model (*SM4. Table S6*), a Spearman correlation analysis showed that the entropy and moiety flux solution and the INCA model flux solution had a relatively good correlation, however, the FBA flux solution had a poor correlation to any other flux solution (Figure 11**.B**).

**Figure 11.**
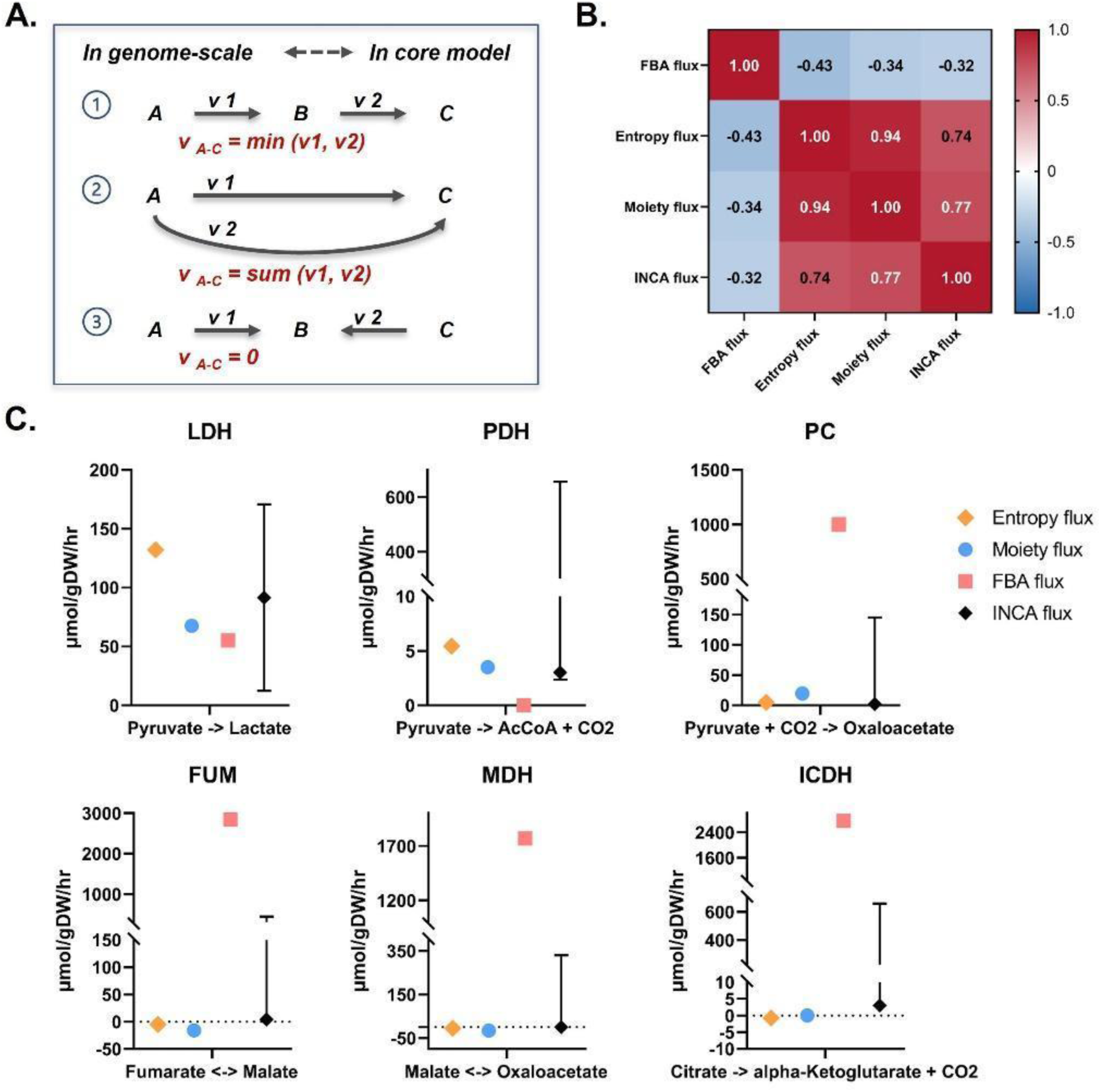
Flux comparison between FBA flux, entropy flux, moiety flux, and INCA model flux within the same central carbon metabolism network for iPSC-derived midbrain-specific dopaminergic neurons. A. Illustration on mapping reaction fluxes at the genome scale to the corresponding reaction flux in the core model. B. Spearman correlation analysis between FBA flux, entropy flux, moiety flux, and INCA flux within the same network. C. Comparison of individual reaction fluxes from glycolysis and tricarboxylic acid cycle.

In terms of the reaction directionality of the nutrient exchange, the entropy, moiety and INCA model flux solutions all showed the uptake of glucose, glutamate, glutamine, and the secretion of lactate and pyruvate in neurons, whereas the FBA flux solution showed the uptake of lactate and pyruvate and no uptake of glutamine. A comparison of internal reactions was then only focused on entropy, moiety, and INCA model flux solutions. Within the subsystem of glycolysis/gluconeogenesis, all relevant reactions share the same directionality between the three flux solutions. The flux value of the reaction converting pyruvate to lactate seen in the entropy and moiety flux solution fell within the INCA flux range. Several reactions of the pentose phosphate pathway showed in opposite directions to INCA model flux solutions, especially for moiety flux solution. They all, however, had low flux compared to glycolytic reactions. Generally, pyruvate can be converted into acetylCoA via pyruvate dehydrogenase (PDH) and join the tricarboxylic acid cycle. Besides, pyruvate anaplerosis can covert pyruvate directly into oxaloacetate via pyruvate carboxylase (PC) to compensate for metabolite loss from tricarboxylic acid cycle due to biomass production [56]. Interestingly, the reaction flux of PDH and PC observed in the moiety flux solution both fell within the INCA flux range. Similar results can be also found in a few other reactions. Regardless of this, three reactions of the tricarboxylic acid cycle showed in opposite directions to INCA model flux solutions, as shown in Figure 11**.C**. A final steady-state flux distribution map based on entropy, moiety and INCA model flux solutions can be seen in Figure 12.

**Figure 12.**
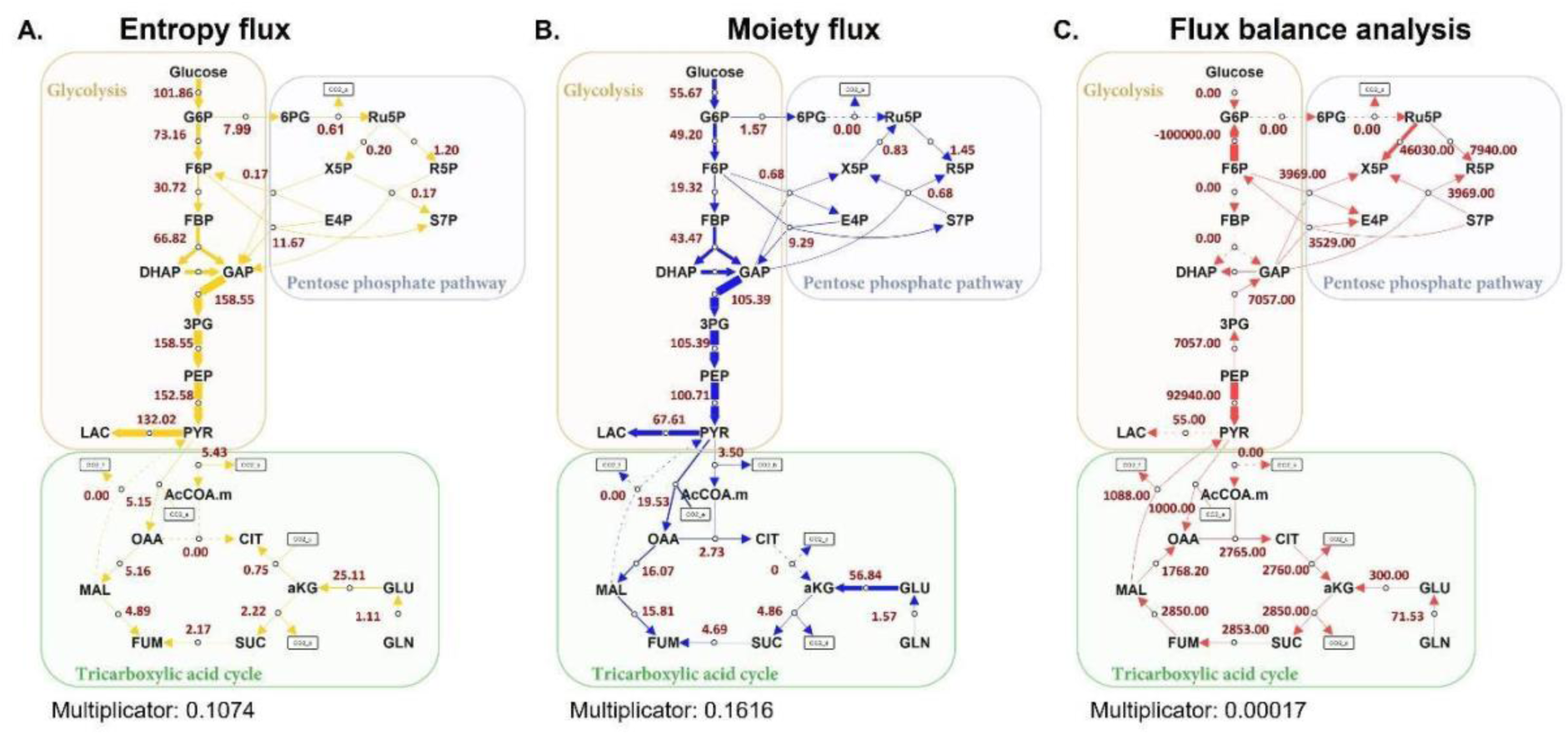
Genome-scale metabolic flux analysis of differentiated dopaminergic neurons projected within central carbon metabolism network. A steady-state flux distribution map over glycolysis, tricarboxylic acid cycle, and pentose phosphate pathway for iPSC-derived midbrain-specific dopaminergic neurons based on entropy (A, shown in yellow), moiety (B, shown in blue), FBA (C, shown in red) flux solutions. Multiplicator represents a multiplicative scaling factor. This factor converts each flux value into a standardized value that enables the use of a uniform arrow thickness scale across all maps, which helps metabolic flux patterns comparison across all maps. The flux unit is μmol/gDW/hr.

Besides, the top 15 active pathways within the genome-scale network based on entropic flux balance analysis were summarised in Figure 13. In comparison to the results of entropy flux, moiety flux solutions exhibited comparable average flux results in most of the subsystems. Higher flux activity can be found in the subsystems of fructose and mannose metabolism, oxidative phosphorylation, glutamate metabolism, and citric acid cycle, whereas lower flux activity was observed in the subsystem of glycolysis/gluconeogenesis.

**Figure 13.**
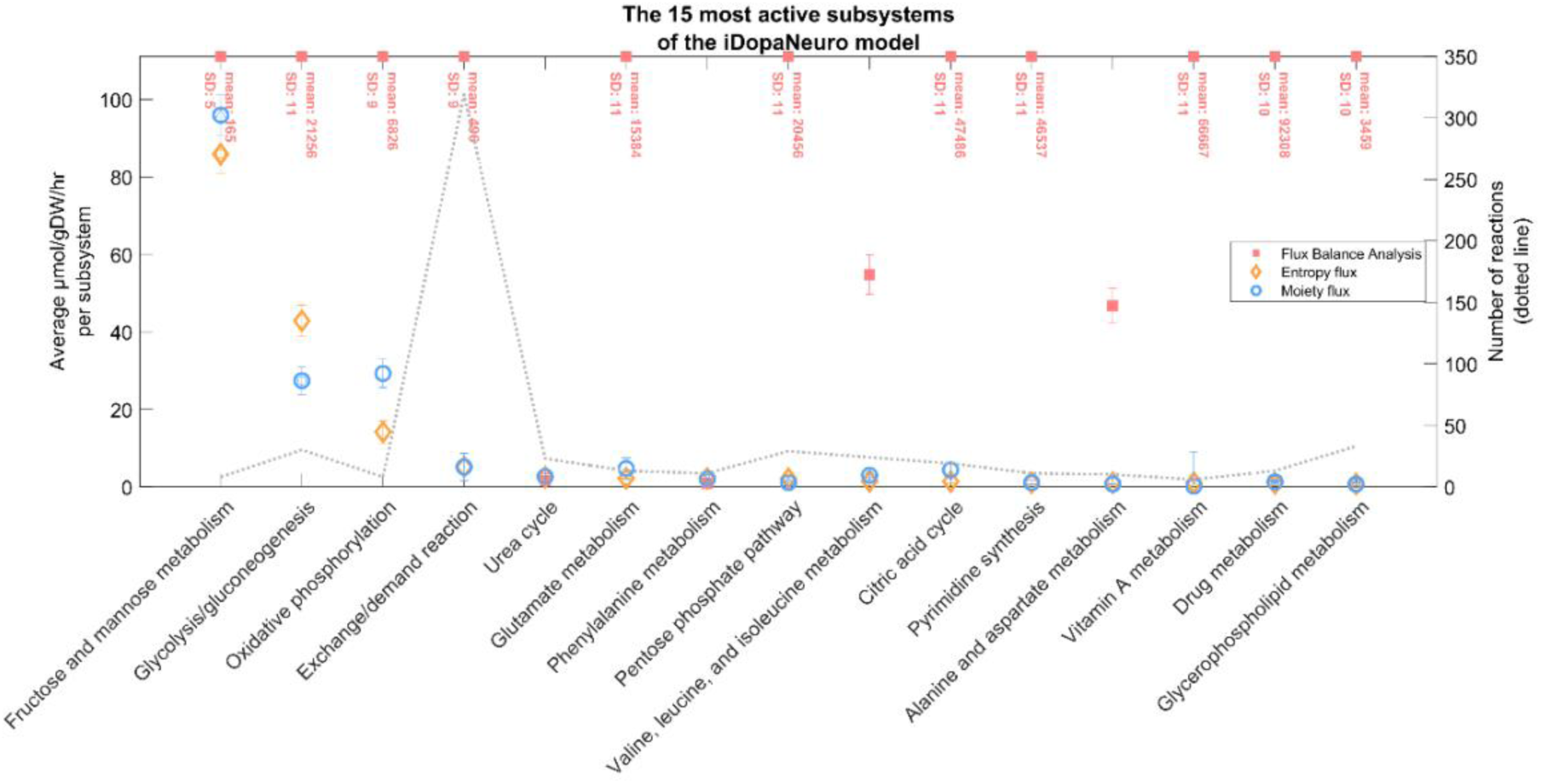
Subsystem activity. Summary of top 15 active pathway subsystems in the iDopaNeuroC model based on the objective function of entropic flux balance analysis, and the correlated results with FBA flux (red), and moiety flux (blue) analysis.

Regarding the phenotypes for carbon source and energy production, entropy and moiety flux shared some similarities and showed a big difference from FBA flux. Notably, with moiety flux, the reaction of ATP synthase in mitochondria produced the most ATP, whereas phosphoglycerate kinase in the cytoplasm produced the most ATP with entropy flux. Both methods relied on glucose as their primary carbon source. (**Table 4**)

**Table 4.** Modelling phenotypes. Comparison of energetic phenotypes obtained using various genome-scale modelling approaches.

|  | FBA flux | Entropy flux | Moiety flux |
| --- | --- | --- | --- |
| Carbon sources |  |  |  |
| Metabolite#1 | Bicarbonate | D-Glucose | D-Glucose |
| Flux | -4268 $\mu\text{mol/gDW/hr}$ | -427.44 $\mu\text{mol/gDW/hr}$ | -427.44 $\mu\text{mol/gDW/hr}$ |
| Formula | $\text{CHO}_3$ | $\text{C}_6\text{H}_{12}\text{O}_6$ | $\text{C}_6\text{H}_{12}\text{O}_6$ |
| Metabolite#2 | D-Glucose | L-Arginine | L-Arginine |
| Flux | -440.40 $\mu\text{mol/gDW/hr}$ | -29.35 $\mu\text{mol/gDW/hr}$ | -27.77 $\mu\text{mol/gDW/hr}$ |
| Formula | $\text{C}_6\text{H}_{12}\text{O}_6$ | $\text{C}_6\text{H}_{15}\text{N}_4\text{O}_2$ | $\text{C}_6\text{H}_{15}\text{N}_4\text{O}_2$ |
| Metabolite#3 | L-Threonine | Glycine | Glycine |
| Flux | -117.30 $\mu\text{mol/gDW/hr}$ | -25.81 $\mu\text{mol/gDW/hr}$ | -27.06 $\mu\text{mol/gDW/hr}$ |
| Formula | $\text{C}_4\text{H}_9\text{NO}_3$ | $\text{C}_2\text{H}_5\text{NO}_2$ | $\text{C}_2\text{H}_5\text{NO}_2$ |
| ATP production |  |  |  |
| 1st ATP source | Nucleotide interconversion | Phosphoglycerate Kinase | ATP synthase |
| Flux | -1e105 $\mu\text{mol/gDW/hr}$ | -158.55 $\mu\text{mol/gDW/hr}$ | 110.88 $\mu\text{mol/gDW/hr}$ |
| 2nd ATP source | Succinate- Coenzyme A Ligase | ATP synthase | Phosphoglycerate Kinase |
| Flux | -1e105 $\mu\text{mol/gDW/hr}$ | 52.23 $\mu\text{mol/gDW/hr}$ | -105.392 $\mu\text{mol/gDW/hr}$ |
| 3rd ATP source | ATP synthase | Pyruvate Kinase | Pyruvate Kinase |
| Flux | 33370 $\mu\text{mol/gDW/hr}$ | 48.15 $\mu\text{mol/gDW/hr}$ | -34.184 $\mu\text{mol/gDW/hr}$ |

## 5. Identification of conserved moieties and new tracer design

A list of 10 conserved moieties within the atomically resolved iDopaNeuroC model was identified (*SM4. Table S7*). Each conserved moiety corresponds to its own subnetwork [27]. To predict the best conserved moieties to label in a future tracer-based metabolomic experiment, we first compared the coverage of internal reactions in every subnetwork. Conserved moiety #59 had the highest coverage of carrying internal reactions flux. Moreover, four of moiety- relevant subsystems were confirmed as top-15 active subsystems in entropy flux-based solution, and two of moiety-relevant subsystems were confirmed as top-15 active subsystems in moiety flux-based solution. Conserved moieties #5 and #56 showed about 50% internal reaction coverage. Nevertheless, they had no active subsystems hits and conserved moieties #56 was not found within any uptaken metabolite in the whole subnetwork. Overall, the conserved moiety #59 showed the most promising candidate because of its subnetwork activity. To further trace that conserved moiety within network through experiments, a partially ^13^C4, D, ^15^N2, ^18^O-labelled thymidine can be designed as a new culture tracer (Figure 14).

**Figure 14.**
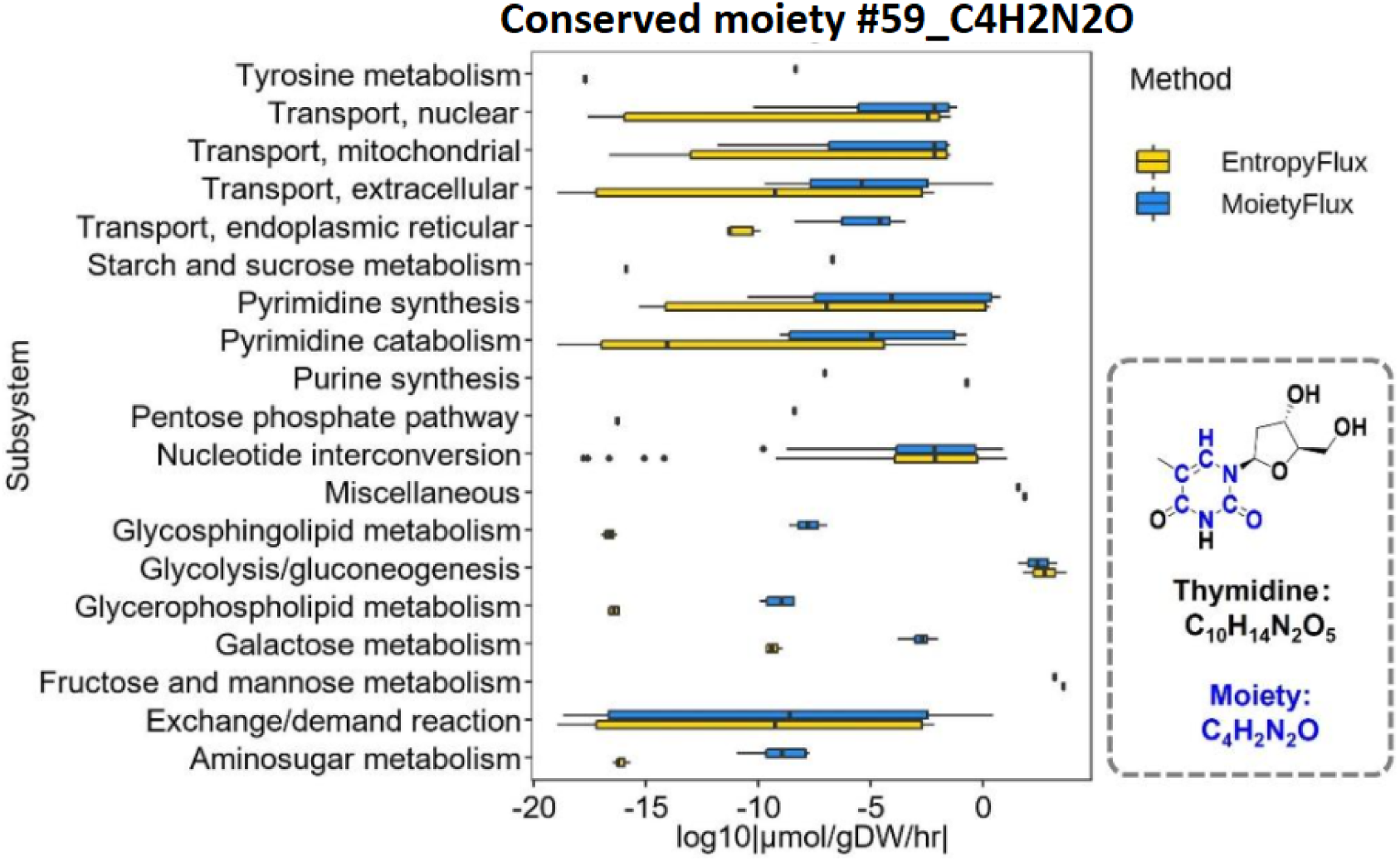
Conserved moiety #59_C_4_H_2_N_2_O. Overall reaction flux plot is classified by metabolic subsystem based on entropy flux and moiety flux predictions. Inside the dotted line illustrates a determined tracer that can be utilised in a new tracer-based metabolomic experiment.

## Discussion

This work describes a COBRA Toolbox v3.4 [13] extension termed fluxTrAM, which aims to standardise the procedure of preparing an atomically resolved metabolic model and processing tracer-based metabolomics data to perform metabolic flux analysis at genome-scale.

### Tracer-based metabolomic data processing

One part of fluxTrAM pipeline is focused on the standardised and high-throughput processing for stable isotope labelling data and ^13^C metabolic flux analysis. The accurate mass isotopologue distribution of the target metabolites depends on good peak detection and extraction. Based on our evaluation, mzMatch-ISO showed higher accuracy than X^13^CMS in collecting targeted isotopologue information. X13CMS is a package mainly designed for untargeted metabolomics [6], and mzMatch is a package for both untargeted and targeted metabolomics, mzMatch-ISO especially showed its unique advantage in isotopic labelling data analysis with a list of target analytes. The function of ***peakDetector*** based on the mzMatch-ISO package presented good consistency in the isotopologue faction analysis for essential amino acids between the labelled and unlabelled groups while showing significant differences for non-essential amino acids. The automated connection between different packages guaranteed that no additional data format manipulation occurs, leading to a consecutive data processing workflow. This was demonstrated in the analysis of a time-course dataset from the formal study set. In addition, the pipeline is flexible in that it allows self-defined parameter settings, and the output result for each function step can be freely exported. It is currently only applicable to processing LC-MS datasets due to the restriction of selected automated peak extraction package. When appropriate packages are developed by the metabolomics community, future development can be extended to LC-MS/MS and GC-MS datasets.

In the application of ^13^C metabolic flux analysis for iPSC-derived dopaminergic neurons, reaction flux was obtained within a central carbon metabolism network model in INCA software. The experimental flux range within 95% confidence interval is valuable in reflecting the basic energy metabolism regulation on a healthy neuronal model. The net reaction flux map showed a very active glycolysis activity compared to the tricarboxylic acid (TCA) cycle and the pentose phosphate pathway. However, this model was built based on many assumptions to obtain a successful fit. Only a few nutrient transports (glucose, glutamine, glutamate) were described in the INCA model compared to 320 exchange reactions in the iDopaNeuroC model. Oxidative sources of unlabelled carbon such as amino acids and fatty acids in the TCA cycle were not considered in this model. The output carbon flux was limited to the secretion of lactate and pyruvate. Besides, metabolites were assigned in one single pool without a compartmental difference. Although the experimental flux solution was based on limited network size and many pre-defined assumptions, it still offers a flux solution that can indicate a feasible cellular activity with statistically approved goodness-of-fit and biological significance.

### Metabolite structure and atom mapping databases

Obtaining metabolite identifiers is critical for expanding the database’s coverage. Similarly, having a wide range of sources increases the probability of obtaining the metabolites that best represent those described in the genome-scale model. An InChI represents various metabolite structures depending on the level of detail included in it (see *Figure S1*). The main layer consists of layers 1 (standardised or not), 2 (chemical formula), 3 (atoms connectivity), and 4 (hydrogen atom connectivity), e.g., InChI=1S/C3H7NO2/c1-2(4)3(5)6/h2H,4H2,1H3,(H,5,6), which represents the alanine molecule; layer 5 adds the charges, e.g., adding p-1, representing the alaninate molecule; and finally, layer 6 indicates the stereochemistry, e.g., adding t2-/m0/s, representing the L-alaninate molecule. As the specificity of the metabolites increases, so does the number of atomically balanced reactions in the database. A high quality atomically resolved genome-scale metabolic model may have multiple applications, some of which were explored in this work. Among the applications are the estimation of fluxes in the metabolic network in combination with MID data [28], identifying conserved moieties in the metabolic network to optimise tracer-based experimental design [26,27], to calculate the potential energy in molecules [57] or to identify the amount of enthalpy change associated with a metabolic reaction via bond enthalpies (See supplementary information).

Metabolite structures may be obtained in a variety of ways, including drawing based on the literature using cheminformatic software [24] or obtained from metabolic databases either manually or computationally given various different identifiers. Different approaches exist to differentiate isomers in order to obtain a consistent representation of a metabolite structure from a biological system, such as the CAS Registry Number, a unique numerical identifier that is independent of any chemical nomenclature system and is widely used to identify chemical substances with a maximum capacity of 1 e9 identifiers [58]; or an InChI, representing a chemical structure in a standardised manner, allowing the generation of a unique identifier for a chemical structure [41], while other cheminformatic formats such as SMILES [42] and chemical tables [45], may use different annotations to represent the same molecule. Nevertheless, assigning the correct identifier for a metabolite in a genome-scale model is a difficult task due to the various isomers that a molecule may have, and more work should be done in this area to avoid mismatches. This is the case of the metabolite nicotinamide adenine dinucleotide, which is represented in the iDopaNeuroC model [30] with the identifier CHEBI:15846 from the ChEBI database, whereas the consistent identifier based on the number of hydrogen atoms should be CHEBI:57540. Tools for mapping identifiers between different biological databases [59], such as BridgeDB [60], the chemical translation service [59,61] and MetaboAnnotator [62], can be useful to assigning metabolite identifiers in accordance with the genome-scale model.

### Integrative metabolic flux analysis and new experimental tracer design

FBA was used to predict the steady-state flux distribution assuming maximization of neuronal ATP production. Entropic flux balance analysis does not maximise the flux of a single reaction but rather assumes the objective function is to maximise the entropy of every forward and reverse internal reaction flux and also predicts a thermodynamically feasible flux vector. In addition, the moiety fluxomics predicts a new genome-scale flux solution in the atomically resolved iDopaNeuroC model by integrating the labelled MID data as extra constraints. The latter two approaches outperformed the FBA method as evaluated by their higher correlations with the flux inference using conventional metabolic flux analysis with INCA while using a central carbon metabolism network model. Moiety flux inference [28] shares some similarities with the entropic flux balance analysis prediction in terms of metabolic phenotype. For instance, glucose was predicted as one of the primary carbon sources for dopaminergic neuron. Nevertheless, it also shows some inconsistencies with entropy modelling solution. Moiety flux predicted a very active ATP production via ATP synthase (four protons for one ATP) in mitochondria, while entropy flux predicted a major ATP production from cytoplasm. While maintaining a comparable ATP production, incorporating labelled MID data reformed some active subsystems of the neuronal metabolism, shown as more active oxidative phosphorylation, glutamate metabolism, and citric acid cycle, and less active glycolysis/gluconeogenesis.

Tracer choice plays an important role in determining the precision of network flux estimation [63,64]. We applied U-^13^C6-glucose as the tracer for iPSC-derived dopaminergic neurons, which is not an ideal tracer to elucidate flux through the pentose phosphate pathway. Although glucose is a major nutrient used for energy production, neurons can utilise multiple other nutrient sources as well. To improve the accuracy of genome-scale flux prediction, a validation experiment with a new tracer is further required. Identification of conserved moieties has strong potential for use in design of tracer-based metabolomic experiments [65]. By isotopically labelling any single atom in a conserved moiety, one can use the predict the reachable set reactions that could contain that isotopic label.

In our study, the results of the genome-scale flux prediction helped to determine optimal tracers by evaluating flux activity of candidate conserved moieties on intracellular metabolism. This will facilitate future study of particularly significant metabolic pathways in more detail. On the other hand, it also contributes to the systematic interaction between genome-scale modelling and experimental validation towards a more efficient level. The conserved moiety proposed in this study, present in the metabolite thymidine, was found in multiple subsystems making it a potential candidate for the design of novel labelled molecules for a tracer-based metabolomics experiments. Ikuno reported an increased level of thymidine in the plasma of prodromal Parkinson’s disease mouse model, which can cause mitochondrial disorders characterized by mitochondrial DNA (mtDNA) depletion [66]. In Anandagopu *et al*. [67], the distribution of thymidine was found to be decreased in PINK1 mRNA *homo sapiens* sequences, a mutation associated with mitochondrial dysfunction and oxidative stress. Not all of the conserved moieties in the iDopaNeuroC were identified. This was hampered by the absence of metabolite structures for some reactants, e.g., lipids with R groups in the structure, as they precluded the atomic resolution of all reactions in the iDopaNeuroC model. However, it was possible to atom map the majority of internal reactions, which permitted the identification of the majority of conserved moieties [26,27] in the iDopaNeuroC model.

## Conclusions

Unlike a snapshot of metabolic phenotypes reflected by metabolite concentration, metabolic fluxes over reaction networks offer a dynamic perspective for characterising cellular metabolism. Current flux analysis is more restricted to small-scale networks and are rarely performed at a genome-scale level. While recent developments in the moiety fluxomics method have addressed this challenge. To standardise and speed up the implementation of metabolic flux analysis, this study developed a semi-automated pipeline, fluxTrAM, that enables the integration of tracer-based metabolomics data with atomically resolved metabolic networks for metabolic flux analysis at genome scale. The first module of fluxTrAM achieved automated processing of tracer-based LC-MS raw data into standardised MID data. The second module achieved atomically resolving any given genome-scale metabolic model, and the results included a cheminformatic database of standardized and context-specific metabolite structures and atom-mapped reactions. Finally, by integrating neuron-specific MID data into the atomically resolved iDopaNeuroC model using the moiety fluxomics method, we successfully inferred the first genome-scale flux solution for human dopaminergic neurons. This moiety flux solution showed good correlation with the entropy flux solution and the conventional C13 flux solution obtained based on central carbon metabolism networks. Thus, global flux monitoring makes it possible to broaden the analysis of metabolic networks beyond central carbon metabolism. It will play an important role in identifying and interpreting vital metabolic dysregulation under diseased conditions for future study. By combining the genome-scale flux solution with conserved moiety analysis, we can also design labelled tracer candidates for new labelling experiments. In this way, we will be able to further validate and advance our understanding of neuronal metabolic regulation in response to any perturbations. This procedure will benefit significantly from the standardisation and high throughput of our proposed fluxTrAM pipeline.

## Data Availability

The fluxTrAM pipeline can be found in the COBRA Toolbox v3.4 [13] (https://github.com/opencobra/COBRA.tutorials/tree/master/analysis/tracerMetabolomics2atomResolvedModels).

The metabolite structures and atom-mapped reactions generated by the fluxTrAM pipeline from the genome-scale models iDopaNeuroC [29], *E. coli* core [47], AGORA2 [54] and Recon3D reconstruction [45] can be found on GitHub (https://github.com/opencobra/COBRA.papers/2023_tracerMetabolomics2atomResolvedModels) and in the VMH database [37].

Scripts for running flux comparison analysis can be found on GitHub (https://github.com/opencobra/COBRA.papers/2023_tracerMetabolomics2atomResolvedModels)

## Acknowledgments

This project has received funding from the European Union’s Horizon 2020 research and innovation programme under grant agreement number (668738, SysMedPD), the European Union’s Horizon Europe Framework Programme (101080997, Recon4IMD) the China scholarship council [No.201806210057], the Dutch National Institutes of Health (ZonMw) TKI-LSH Neuromet project (LSHM18092) and the Dutch Research Council (NWO)

’Investment Grant NWO Large’ program, for the ’Building the infrastructure for Exposome research: Exposome-Scan’ project (No. 175.2019.032).

## Author Contributions

Luojiao Huang: Conceptualisation, Methodology, Software, Validation, Formal analysis, Writing - Original Draft & Visualisation.

German Preciat: Conceptualisation, Methodology, Software, Validation, Formal analysis, Writing - Original Draft & Visualisation.

Jesus Alarcon: Conceptualisation, Methodology, Software & Visualisation. Edinson L. Moreno: Methodology.

Agnieszka Wegrzyn: Supervision, Writing - Review & Editing. Ines Thiele: Methodology & Supervision.

Emma Schymanski: Methodology & Supervision.

Amy Harms: Methodology & Supervision, Writing - Review & Editing.

Ronan M.T. Fleming: Conceptualisation, Software, Formal analysis, Writing - Review & Editing, Supervision, Project administration & Funding acquisition.

Thomas Hankemeier: Writing - Review & Editing, Supervision, Project administration & Funding acquisition.

All authors have read and agreed to the published version of the manuscript.

