## Supplementary Information for "fluxTrAM: Integration of tracer-based metabolomics data into atomically resolved genome-scale metabolic networks for metabolic flux analysis"

### Supplementary Materials

#### 1. Inconsistencies in atom mapping data

Not all of the metabolites from the HMDB database were collected, resulting in a low collection of metabolites. This was due to connection problems with the database, as shown by the The HyperText Transfer Protocol (HTTP) status 503, which indicates that the server is not ready to handle the request.

Inconsistent metabolites or reactions can cause unbalanced reactions. With the iDopaNeuroC model, 15 metabolites were always present in at least one of the 100 unbalanced reactions; of these metabolites, 9 have non-chemical atoms that are used to represent a variable metabolite pool or to highlight the reacting zone. Additionally, there are inconsistencies between the information from the sources and the information in the iDopaNeuroC model in 11 metabolites, such as metabolite 9E-elaidic acid, where the InChI with the highest score has a different chemical formula than the formula described in the iDopaNeuroC model, C18H34O2 in all the sources and in the model $C_{39}H_{64}N_7O_{17}P_3S$ . Furthermore, because the metabolites were obtained from open databases, the model's identifiers in some cases represent a molecule with no stereochemistry or charge, as demonstrated by the metabolite L-arginine, from which the VMH alone obtained the highest score. L-arginine is represented in on the VMH [31] as arg_L, in KEGG [39] as C00062 in PubChem [43] as 6322 , in HMDB [40] as HMDB00517 and in ChEBI [44] as CHEBI:16467.

##### 1.1 Inconsistencies caused by R groups

The R groups can cause inconsistencies because they represent a substructure that, because they are not defined, can result in unbalanced reactions, such as reaction sphingomyelin phosphodiesterase (VMH ID: HMR_0795).

| Reaction | VMH ID | Formula |
|---|---|---|
| Glucosylceramidase | GBA | h2o[c] + gluside_hs[c] -> glc_D[c] + crm_hs[c] |
| Glucosylceramidase (lysosome) | GBAl | h2o[l] + gluside_hs[l] -> crm_hs[l] + glc_D[l] |
| S-Adenosyl-L-Methionine: Phosphatidylethanolamine N-Methyltransferase | HMR_0653 | amet[c] + pe_hs[c] -> 2 h[c] + ahcys[c] + M02686[c] |
| Ceramide glucosyltransferase | HMR_0761 | crm_hs[c] + udpg[c] -> h[c] + udp[c] + gluside_hs[c] |
| Sphingomyelin phosphodiesterase | HMR_0795 | h2o[c] + sphmyln_hs[c] -> h[c] + crm_hs[c] + cholp[c] |
| Phosphatidic acid phosphatase | PPAP | h2o[c] + pa_hs[c] -> pi[c] + dag_hs[c] |
| Phosphatidylserine decarboxylase | PSDm_hs | h[m] + ps_hs[m] -> co2[m] + pe_hs[m] |
| Phosphatidylserine synthase | PSSA1_hs | ser_L[c] + pchol_hs[c] <=> chol[c] + ps_hs[c] |
| Phosphatidylserine synthase | PSSA2_hs | ser_L[c] + pe_hs[c] <=> etha[c] + ps_hs[c] |
| Psdm_Hsc | PSDm_hsc | h[c] + ps_hs[c] -> co2[c] + pe_hs[c] |
| Sphingomyelin synthase | SMS | pchol_hs[c] + crm_hs[c] -> dag_hs[c] + sphmyln_hs[c] |

##### 1.2 Inconsistencies caused by protons

Inconsistencies with protons are caused by metabolite pH, which changes the number of hydrogens, resulting in unbalanced reactions, such as reaction N-Acyl-Aliphatic-L-Amino Acid Amidohydrolase (VMH ID: RE2642C)

| Reaction | VMH ID | Formula |
|---|---|---|
| Cardiolipin synthase | CLS_hs | cdpdag_hs[c] + pglyc_hs[c] -> h[c] + cmp[c] + clpn_hs[c] |
| Glucosylceramidase | GBA | h2o[c] + gluside_hs[c] -> glc_D[c] + crm_hs[c] |
| Glucosylceramidase (lysosome) | GBAl | h2o[l] + gluside_hs[l] -> crm_hs[l] + glc_D[l] |
| S-adenosyl-L-methionine: phosphatidyl-N-dimethylethanolamine N-methyltransferase | HMR_0657 | amet[c] + M02758[c] -> ahcys[c] + pchol_hs[c] |
| Ceramide glucosyltransferase | HMR_0761 | crm_hs[c] + udpg[c] -> h[c] + udp[c] + gluside_hs[c] |
| Metabolism of LeuSerTrp (formation/degradation) | LEUSERTRPr | 2 h2o[c] + leusertrp[c] <=> ser_L[c] + leu_L[c] + trp_L[c] |
| Lysophospholipase | LPASE | h2o[c] + lpchol_hs[c] -> h[c] + Rtotal[c] + g3pc[c] |
| Nicotinate D-ribonucleoside kinase | NICRNS | atp[c] + nicrns[c] -> h[c] + adp[c] + nicrnt[c] |
| Nucleotide phosphatase | NP1 | h[c] + nac[c] + r1p[c] -> pi[c] + nicrns[c] |
| Phospholipase A2 | PLA2_2 | h2o[c] + pchol_hs[c] -> h[c] + Rtotal2[c] + lpchol_hs[c] |
| Phospholipase A2 (extracellular) | PLA2_2e | h2o[e] + pchol_hs[e] -> h[e] + lpchol_hs[e] + Rtotal2[e] |
| RE1266C | RE1266C | o2[c] + 4mop[c] -> h[c] + co2[c] + CE2028[c] |
| Aminoacid N-acetyltransferase | RE2031M | accoa[m] + ala_L[m] <=> h[m] + coa[m] + CE1554[m] |
| N-acyl-aliphatic-L-amino acid amidohydrolase | RE2642C | h2o[c] + CE1554[c] <=> ac[c] + ala_L[c] |

#### 1.3 Missing reactions

Finally, three of the missing reactions have stoichiometry that cannot be represented with integers, resulting in the failure to generate an MDL RXN reaction. Additionally, not all metabolites were present in the degradation (VMH id: HDL_HSDEG) and formation (VMH id: HDL_HSSYN) of HDL due to the limited information available about metabolites in the iDopaNeuroC model.

| Reaction | VMH ID | Formula |
|---|---|---|
| Cytochrome C oxidase; mitochondrial complex IV | CYOOm3 | o2[m] + 7.92 h[m] + 4 focytC[m] -> 1.96 h2o[m] + 4 h[c] + 4 ficytC[m] + 0.02 o2s[m] |
| Degradation of HDL | HDL_HSDEG | h2o[e] + hdl_hs[e] -> 2 chsterol[e] + 2 pchol_hs[e] + Rtotal[e] + Rtotal2[e] + Rtotal3[e] + glyc[e] + HC00004[e] + HC00006[e] + HC00007[e] + HC00008[e] + HC00009[e] |
| Formation of HDL | HDL_HSSYN | 2 chsterol[e] + 2 pchol_hs[e] + tag_hs[e] + HC00004[e] + HC00006[e] + HC00007[e] + HC00008[e] + HC00009[e] -> hdl_hs[e] |
| HMR_0017 | HMR_0017 | M02909[e] -> 0.125 octa[e] + 0.125 dca[e] + 0.125 but[e] + 0.125 C01601[e] + 0.125 M02108[e] + 0.125 M03117[e] + 0.125 M03134[e] + 0.125 caproic[e] |
| Phosphate transport via Na+ symporter | PIt8 | 1.5 na1[e] + pi[e] <=> pi[c] + 1.5 na1[c] |

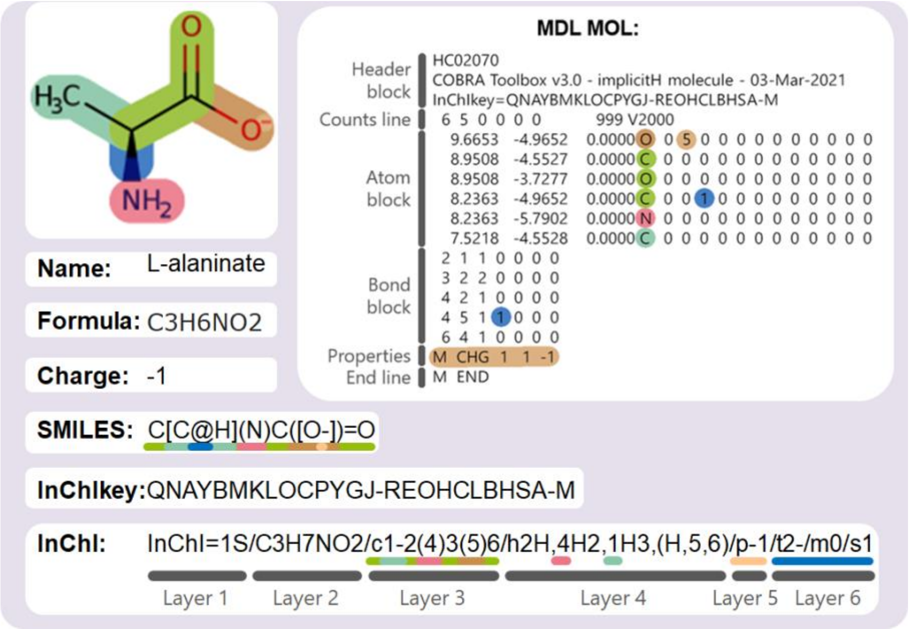


**Figure S1. Comparison of metabolite structure formats.**

An L-alaninate molecule represented by a hydrogen-suppressed molecular graph (implicit hydrogens). The main branch of the molecule can be seen in green; the additional branches can be seen in brown, pink and turquoise. The stereochemistry of the molecule is highlighted in blue, the double bond in dark green and the charges in light brown. The same colours are used to indicate where this information is represented in the different metabolite strucure formats. The InChI is divided into layers, each of which begins with a lowercase letter, except for Layers 1 and 2. Layer 1 indicates if the InChI is standardised. Layer 2 shows the chemical formula in a neutral state. Layer 3 indicates the connectivity between the atoms (ignoring hydrogen atoms). Layer 4 demonstrates the connectivity of the hydrogen atom. Layer 5 indicates the charge of the molecule. Layer 6 shows the stereochemistry. Additional layers can be added, but they cannot be represented with a standard InChI.

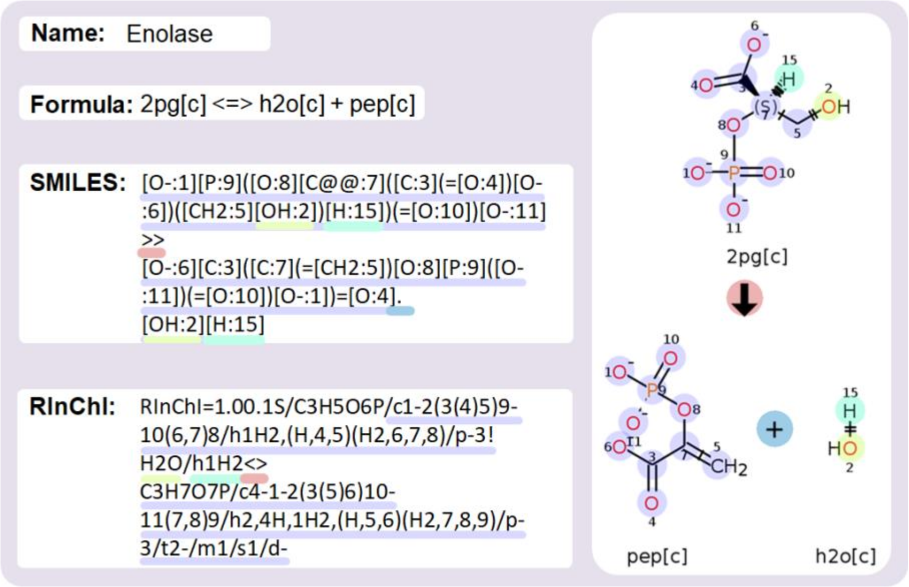


**Figure S2. Chemoinformatic formats for chemical reactions.**

Atom-mapped representation of enolase reaction in different chemoinformatic formats. The larger moiety in the reaction is shown in blue, the hydroxide moiety in yellow, the hydrogen atom in green, the sum sign in dark blue and the reaction sign in red.
